# Longitudinal consensus clustering reveals the functional architecture of the developing rat brain

**DOI:** 10.64898/2026.06.09.731184

**Authors:** Daniel McLoone, Andrew Breen, Andrew Harkin, Clare Kelly

## Abstract

Resting-state fMRI-based mapping of functional networks in developing human populations holds promise for enhancing the diagnosis and treatment of psychiatric disorders, which often emerge during adolescence. Preclinical models offer experimentally tractable lifespans to investigate these trajectories longitudinally, while also enabling direct experimental manipulations that can reveal causal influences on brain and behavioural outcomes. However, unlike in the human neuroimaging field, preclinical neuroimaging often lacks standardisation, making comparison across studies and translation of findings more difficult. In particular, we lack developmentally informed functional brain atlases. Working with a longitudinal sex-balanced resting fMRI dataset collected using a standardised protocol, we aimed to identify robust functional networks in the rat brain and investigate how they change over development (P28, P35, P49, P70 and P91). Employing a consensus clustering approach informed by cluster quality metrics, we mapped functional network development of the rat brain across multiple spatial scales (*k*=3, *k*=5 and *k*=7) from juvenile (pre-puberty) to early adulthood. Force-directed spring embeddings and graph metrics showed global shifts in brain organisation; with increased global system segregation over time, while global network integration (participation coefficient) decreased. At the network-level, the rat brain was characterised by functionally heterogenous network maturation that recapitulated sensorimotor – higher cognitive gradients of development observed in humans. Longitudinal mapping over the adolescent period revealed both linear and non-linear developmental trajectories, and shows that maturation of the rat brain is characterised by network fractionation and delayed cortical specialisation, with the brain transitioning from globally interconnected juvenile networks into functionally segregated adult networks. The functional architecture we describe offers a generalisable structure and nomenclature for the preclinical imaging community, analogous to the Yeo 7-network parcellation in humans. By openly sharing all data, analytic resources, and outputs with the community, this work provides a developmentally informed functional atlas developed under a standardised protocol for anaesthesia and data acquisition and offers resources that will enable preclinical researchers to further advance our understanding of brain development during this critical period.

## 1. Introduction

Using resting state fMRI approaches to map functional networks continues to expand our understanding of the brain’s functional architecture, its development across the lifespan, and the plethora of influences that shape neurobiological and behavioural outcomes. Interrogation of the functional integrity of these networks holds promise for enhancing the diagnosis, treatment and prevention of neurological and psychiatric disorders. In humans, functional neuroimaging has enabled identification of network-based subtypes of major depressive disorder^1^ and autism^2^, common network signatures of psychedelic drugs^3^, as well as targets for deep brain stimulation and transcranial magnetic stimulation^4^.

A developmental perspective is essential to a full understanding of neuropsychopathology, particularly during late childhood and adolescence: a period characterised by physical^5^, cognitive and emotional maturation^6^. At the neural level, drivers of these changes include myelination of axons and synaptic pruning^7–9^, as well as changes in cell morphology, neurotransmission, and receptor density^10^, which begin early in life and extend into adulthood. Consequently, the brain is vulnerable to adverse experiences during this period^11,12^, which can set in train neurodevelopmental trajectories that depart from health and towards disorder^13^.

While longitudinal neuroimaging in large human cohorts has advanced our understanding of trajectories of brain development, these studies require considerable funding and effort, are vulnerable to participant attrition, and take years to complete. This is where preclinical studies can prove particularly valuable, and can complement the human literature on development. Preclinical models, such as rodents, allow direct experimental manipulations (e.g., chronic exposure to stress) that permit causal inferences about the effects of a wide range of influences on brain and behaviour. Because they reach adulthood in approximately 3 months, rodent lifespans offer experimentally tractable timeframes for the acquisition of longitudinal data, which are also essential to a causal understanding. Importantly, preclinical MRI studies can use the same neuroimaging methods routinely applied in humans, providing a translational bridge from the causal and mechanistic insights obtainable only in animal models. Finally, conducting longitudinal neuroimaging in rodents presents distinct methodological advantages. In line with the “3Rs” (Replacement, Reduction, and Refinement), longitudinal designs enable a reduction in animal numbers, thanks to the greater statistical power achievable, relative to cross-sectional designs^14–16^

Despite the strengths of rodent fMRI, the field has not yet received as much research attention as human neuroimaging, particularly concerning development. Crucially, the field lacks standardised and widely applicable functional parcellations of the rat brain. Functional atlases enable the reduction of high-dimensional voxel data into manageable networks of interest, reducing complexity, increasing statistical power, and providing a common nomenclature for analysis and interpretation across studies^17,18^, and importantly, across species. In humans, functionally derived networks such as the Yeo 7-network and 17-network parcellations^19^ provide a generalisable, interpretable framework for data analysis. In contrast, rat fMRI studies often rely on anatomical atlases^20–22^. While anatomical atlases provide excellent spatial topography, they do not group regions by functional homogeneity. Although several rat brain functional parcellations exist, their application is limited by methodological factors, such as the inclusion of male animals only and the conditions in which their functional scans were carried under (e.g., isoflurane only sedation^20^ or awake state^23^). Recently, the field has recognised the need for standardisation of preclinical fMRI acquisition protocols to aid in the dissemination, aggregation and reuse of data^24,25^. A collaborative initiative identified and validated a standardised sedation protocol using a combination of low dose isoflurane and medetomidine, which preserves functional connectivity^24^. Subsequently, there is a need for a corresponding functional atlas that captures network architecture under these standardised conditions.

To address this gap, this study aimed to define a robust, generalisable set of functional rat brain networks and map their assembly across development, in a publicly shared longitudinal, sex-balanced developmental dataset collected using a standardised protocol with five developmental timepoints spanning from juvenile to early adulthood. Consistent with observations in humans, we hypothesised that functional networks would exhibit increased segregation alongside a decrease in non-specific network connectivity over development, and that higher-order networks would mature later than sensorimotor networks. To balance anatomical specificity with functional generalisability, we anchored our functional data to the updated Waxholm Space anatomical atlas^26^, enabling cross-modal integration.

## 2. Methods

Extensive methodological details for this dataset are shared in a companion paper^27^; details relevant to the current study are reproduced below. All imaging data are publicly available on OpenNeuro ds007859; https://openneuro.org/datasets/ds007859/; all code is available at https://codeberg.org/mclooned/rat_developmental_clustering.

### 2.1 Animals

All animal experiments and procedures were conducted in compliance with the European Directive 2010/63/EU on the protection of animals used for scientific purposes, approved by the Animal Research Ethics Committee in Trinity College Dublin and carried out under licence granted by the Health Products Regulatory Authority (project authorisation AE19136/P173).

Male and female Wistar Han rats were obtained from Inotiv (United Kingdom) and were bred in house. Rats were weaned on postnatal day 21 (P21), with the day of birth considered P0. Animals were housed in pairs in individually ventilated cages, with nesting material, a red polycarbonate tunnel, a red tall rat loft, and ad libitum access to food and water. Animals were kept in climate-controlled rooms set to 22°C (±2°C) with relative humidity levels of 50% and a 12-hour light-dark cycle. Two male experimenters handled and carried out all experimental procedures on the rats, while staff providing general animal care (water, bedding, etc.) were both male and female.

### 2.2 MRI acquisition

MRI scanning was performed at approximately age P28, P35, P49, P70 and P91 (**Figure 1**). All imaging was performed on a 7T Bruker BioSpec 70/30 USR scanner equipped with a 1H receive-only 8 x 1 rat surface array coil, 72mm transmit coil, BGA9SHP gradients and Paravision 7 software. Animal preparation, sedation, and image acquisition were conducted in accordance with the StandardRat consensus protocol^24^. Briefly, animals were maintained under a combined isoflurane and medetomidine anaesthesia regime with continuous physiological monitoring. Functional scans were acquired using a gradient-echo EPI sequence (TR = 1000 ms; TE = 17 ms; in-plane resolution = 0.4 × 0.4 mm; slice thickness = 1 mm). This was paired with a structural T2-weighted turboRARE sequence for anatomical reference (TR = 2500 ms; TE = 30 ms; in-plane resolution = 0.2 × 0.2 mm; slice thickness = 1 mm). Detailed information regarding the anaesthesia induction timeline, hardware specifications, and sequence parameters have been described previously in the original dataset publication^27^.

**Figure 1.**
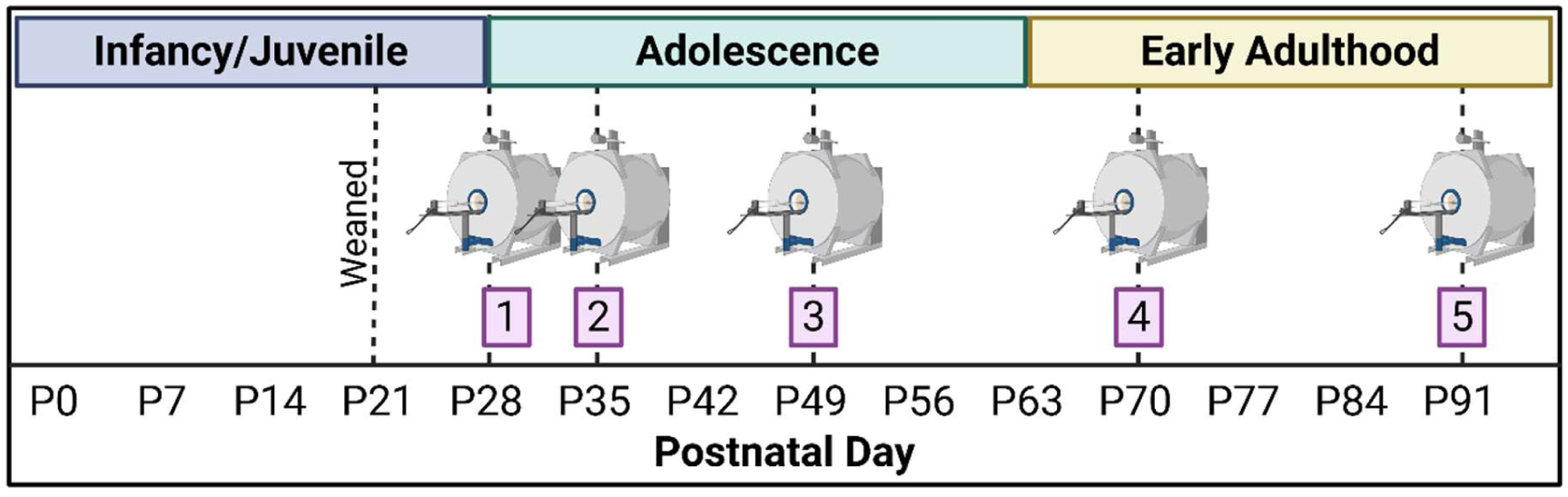
Timeline of functional scan acquisition. Animals were scanned at approximately P28, P35, P49, P70 and P91. Created in BioRender. Mc Loone, D. (2026) https://BioRender.com/ pc7gguq

### 2.3 Data processing

All neuroimaging data was structured in BIDS format and preprocessed using a combination of AFNI^28^, ANTs^29^, and RABIES preprocessing software^30^. Comprehensive details regarding the pipeline, including environment specifications, are available in the dataset publication^27^.

Briefly, to ensure accurate normalisation across the developmental lifespan of the cohort, a study-specific unbiased population-average anatomical template was generated using optimized_antsMultivariateTemplateConstruction (https://github.com/CoBrALab/optimized_antsMultivariateTemplateConstruction)^31^, with initial masks generated using brain extraction net (BEN)^32^. This template was subsequently registered to the SIGMA in vivo rat atlas^20^.

Functional data was preprocessed using Rodent Automated Bold Improvement of EPI Sequences (RABIES)^30^. Timeseries underwent susceptibility distortion correction and non-linear registration to the study-specific template. Confound correction was performed on the native space EPI timeseries. This included voxelwise detrending, bandpass filtering (0.01–0.1 Hz, with 30 s edge-artefact removal from each side of the timeseries), and the regression of six rigid-body motion parameters alongside the global signal. The data was spatially smoothed using a 0.5 mm full-width at half maximum (FWHM) Gaussian filter. Following visual quality control, 11 scans were excluded due to preprocessing failures, resulting in a final sample of 168 functional scans for the current analysis: P28: n = 27; P35: n = 33; P49: n = 36; P70: n = 36; P91: n = 36.

### 2.4 Regions of interest

The brain atlas from the SIGMA Rat Brain Templates and Atlases Version 2.0 which specifies anatomical regions according to the updated Waxholm atlas^26^ was used in this study (https://zenodo.org/records/10635831)^20^. To avoid constraining our clustering algorithm to produce bilateral clusters, potentially masking developmental changes, we created a version of the SIGMA_InVivo_Anatomical_Brain_Atlas.nii.gz atlas that split each of the 222 bilateral ROIs into left and right ROIs, resulting in a total of 444 ROIs. We then registered our population average to the SIGMA_InVivo_Anatomical_Brain_template.nii.gz and applied the inverse transform to obtain the ROIs in our study-specific population average space.

To address areas of signal drop-out and incomplete coverage due to FOV (e.g., in posterior cerebellum, brainstem, and anterior olfactory regions), we created a group-level functional mask by summing together all subject-level study-specific population average space masks, and retaining only those voxels present in all subjects. The study-specific SIGMA ROI template was then multiplied by the resulting inclusive mask. We also removed ROIs with less than 10 voxels (see **supplemental material 1** for included/excluded ROIs). This yielded a lateralised brain atlas in our population space with a total of 292 ROIs. The mean timeseries was then extracted from each ROI (using AFNI’s 3dROIstats -nzmean) and we computed the Pearson correlation between each ROI pair, to construct a 292×292 correlation matrix quantifying functional connectivity, for each subject at each age.

### 2.5 Clustering

#### 2.5.1 Eta-squared

To identify functional networks within the rat brain, we applied a data-driven clustering approach previously applied to human fMRI data^33^. Prior to applying the clustering algorithm, we first quantified the spatial similarity of the whole-brain functional connectivity profiles between each of the 292 regions of interest using the *eta*^2^ statistic. As previously demonstrated by Cohen et al.^34^, the *eta*^2^ statistic provides a more sensitive measure of similarity, since it takes into account differences in scaling and offset between two connectivity maps, which are unaccounted for by correlations.

Using the Pearson whole-brain connectivity vector for each ROI as the input, *eta*^2^ was computed for every ROI pair as follows:

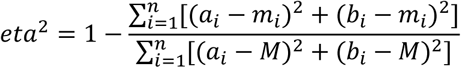

where 𝑎 and 𝑏 are the whole-brain connectivity vectors being compared. 𝑛 is the total number of ROIs in the atlas. The variable 𝑎_i_ is the correlation value describing the functional connectivity between ROI 𝑎 and ROI 𝑖, while 𝑏_i_represents the functional connectivity between ROI 𝑏 and ROI 𝑖. The term 𝑚_i_ is the pairwise mean of 𝑎_i_ and 𝑏_i_. 𝑀 is the grand mean of all functional connectivity values across the vectors for both ROIs 𝑎 and 𝑏 combined. This calculation yields an *eta^2^* matrix representing the profile of similarity across all regions, where values closer to 1 indicate more similar connectivity profiles.

This resulted in a 292 x 292 *eta^2^* similarity matrix for each subject at each age, resulting in 168 individual matrices for subsequent clustering analysis.

#### 2.5.2 Spectral Clustering

As previously described by Kelly et al.^29,31^, we used the spectral clustering toolbox written in *MATLAB* by Verma and Meilă (https://sites.stat.washington.edu/spectral/). Specifically, we employed the Meila-Shi (multicut) algorithm^36^, which first computes the eigen-decomposition of the normalised Laplacian of our *eta^2^* similarity matrix. This process produces eigenvectors, which we then used as the input for the *k*-means clustering algorithm to partition the data into *k* clusters based on the *k* highest-ranking eigenvectors. This spectral clustering algorithm was applied to the eta^2^ matrices computed above to partition the 292 brain ROIs into *k* clusters, where *k* ranged from 2 to 15, where 15 was considered an upper limit on a computationally tractable and interpretable network scheme for the purposes of our study.

#### 2.5.3 Consensus Clustering

The subject-level results generated from spectral clustering above were then aggregated to determine a single, stable group-level solution. First, for each subject (*s*) and each *k*, an individual adjacency matrix was constructed, where each element of the matrix, 𝑎^(s)^= 1 if ROI 𝑖 and ROI 𝑗 are assigned to the same cluster *k*, and 0 otherwise. A group consensus matrix was then constructed for each value of *k* by averaging across these individual adjacency matrices. This 292 × 292 matrix represents the stability of brain network organisation across subjects, where each element 𝐶_ij_ contains the proportion of subjects in which ROI 𝑖 and ROI 𝑗 were assigned to the same cluster. Spectral clustering was then applied to this group consensus matrix to produce the final group-level community assignments for each *k*. This process of spectral and consensus clustering was carried out independently for each age to yield age-specific clustering solutions. Clustering steps are summarised in **Figure 2**. All clustering steps were carried out in *MATLAB* 2025b.

**Figure 2.**
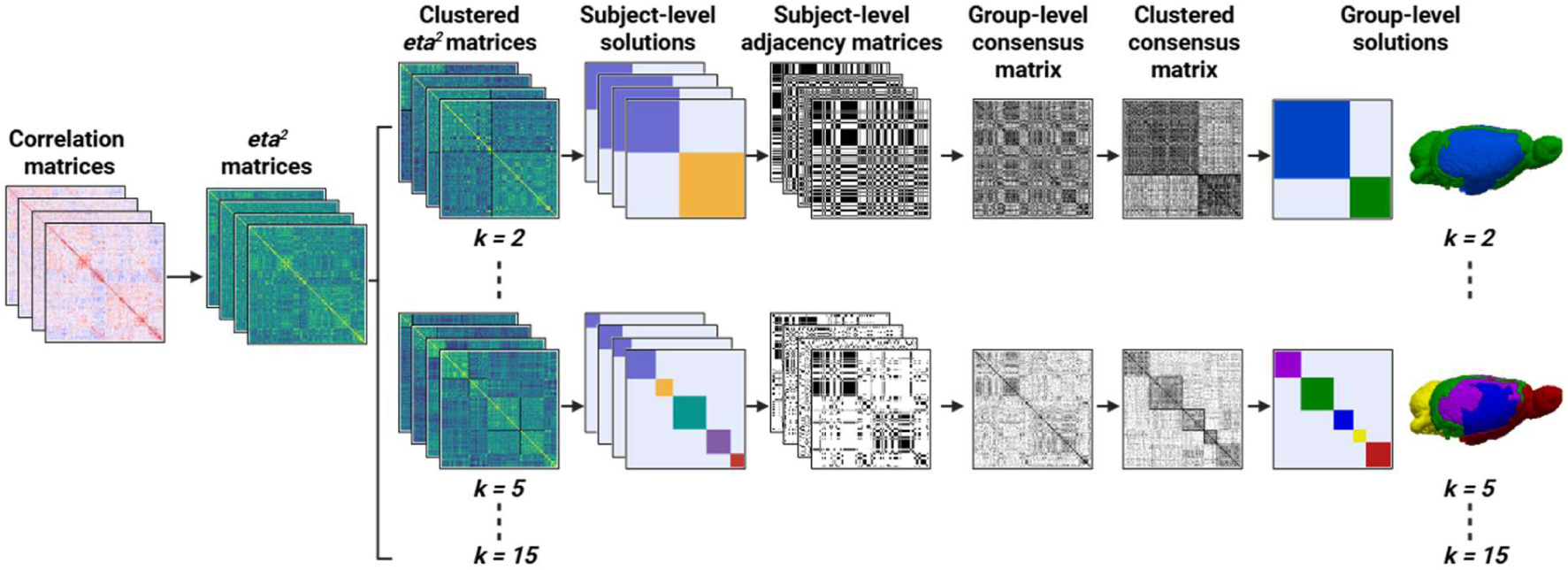
Summary of the steps used to generate group-level consensus clusters. The *eta^2^* statistic for each ROI pair was calculated using the Pearson correlation matrices as the input. Subject-level *eta^2^* matrices were clustered to generate subject-level solutions. From these, an adjacency matrix was created for each subject, with regions clustered together given a value of 1, while regions not clustered together were given a value of 0. A consensus matrix was constructed for each solution of *k* by averaging across all subject-level adjacency matrices. The consensus matrix was then clustered to generate the final group-level solutions for each value of *k.* These steps were applied independently at each age.

### 2.6 Identification of the optimal cluster solution

A fundamental challenge in unsupervised clustering is the selection of the optimal cluster solution (*k*), as *k* is a free parameter and the number of clusters in the rodent brain are unknown. As there is no universally agreed-upon heuristic for identifying the true number of functional networks, we used a combination of metrics. To evaluate the internal validity and separation of each clustering solution, we used three metrics: the modified silhouette score, the Davies-Bouldin index, and mean consensus. To evaluate the robustness and replicability of the clusters across subjects, we assessed three stability metrics: mean variation of information, mean Cramér’s V, and mean percent agreement.

#### 2.6.1 Mean consensus

Mean consensus measures the average probability that regions grouped together in the final consensus solution were clustered together across individual subjects. It is calculated as:

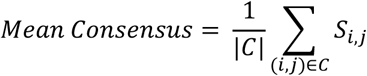

where 𝐶 represents the set of all ROI pairs (𝑖, 𝑗) that are assigned to the same cluster in the final group-level solution, and |𝐶| represents the total number of such intra-cluster pairs. The term 𝑆_i,j_ denotes the specific value from the group consensus matrix (i.e., the proportion of subjects in which region 𝑖 and region 𝑗 were clustered together). A higher value indicates that regions were consistently grouped into the same cluster across subjects.

#### 2.6.2 Modified silhouette

To evaluate cluster cohesion and separation, we computed a modified silhouette score. Rather than calculating a silhouette value for each individual region of interest, the score was computed at the cluster level. Furthermore, because our input matrices represent spatial similarity (𝑒𝑡𝑎^2^) these values were first converted to a measure of distance by subtracting the 𝑒𝑡𝑎^2^values from 1.

For a given cluster 𝐶, the cluster-level silhouette score 𝑆(𝐶) is calculated as:

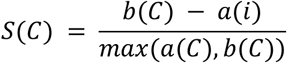

where 𝑎(𝐶) represents the average distance (1 − eta^2^) between all pairs of ROIs within cluster 𝐶. The term 𝑏(𝐶) represents the minimum average distance between the ROIs in cluster 𝐶 and the ROIs in any other single cluster (i.e., the distance to the nearest neighbouring cluster). The resulting score ranges from −1 to 1. A high positive value indicates that the ROIs within the cluster are cohesive and well-separated from other clusters. To evaluate the overall validity of a specific *k* solution, the mean of the cluster-level silhouette scores was calculated.

#### 2.6.3 Davies-Bouldin index

The Davies-Bouldin index evaluates the ratio of “within-cluster scatter” to “between-cluster separation.” For this metric, the 292 x 292 *eta^2^* similarity matrix was used as the input, representing the whole-brain connectivity profile for each region of interest. For a partition with *k* clusters, the Davies-Bouldin index was calculated as follows:

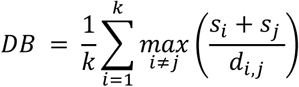

where 𝑠_i_ and 𝑠_j_ represent the intra-cluster scatter for clusters 𝑖 and 𝑗 respectively, computed as the Euclidean sum of squares of the similarity profiles within each cluster. The term 𝑑_i,j_ denotes the inter-cluster separation, computed as the Euclidean distance between the centroids of clusters 𝑖 and 𝑗. For each cluster 𝑖, the metric identifies the neighbouring cluster 𝑗 that produces the maximum ratio (i.e., the most similar, “worst-case” neighbour). The final DB index is the average of these maximum ratios. Consequently, a lower DB value indicates an optimal solution characterised by cohesive clusters that are well-separated in their connectivity profiles.

#### 2.6.4 Variation of information

Variation of Information (VI) is an information-theory measure of the distance between two clustering solutions. To evaluate the stability of a cluster solution we evaluated the current cluster solution (e.g., *k=5*) against every other solution (e.g., *k=2, 3, 4, 6…15*). The VI between a target clustering solution 𝐶_k_ and any other solution 𝐶_kr_ is defined as:

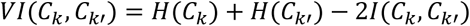

Where 𝐻(𝐶) denotes the entropy of a given clustering solution (representing the uncertainty or information content of the cluster sizes) and 𝐼(𝐶_k_, 𝐶_kr_) represents the mutual information between the two partitions (representing the information shared between them). The Mean VI for solution k was calculated by getting the mean of the pairwise VI distances between 𝐶_k_ and every other solution 𝐶_kr_ (𝑘′ ≠ 𝑘). Because VI represents a true distance metric in information space, lower Mean VI values indicate that the spatial boundaries of the target cluster solution are highly stable and preserved across solutions of *k*.

#### 2.6.5 Cramér’s V

Cramér’s V is a measure of association between two nominal variables, used here to quantify the spatial agreement between the categorical cluster labels of different cluster solutions. Similar to the Mean Variation of Information approach above, we investigated the stability of a target clustering solution by computing its mean Cramér’s V against all other solutions for *k*. For any two cluster solutions being compared, Cramér’s V is calculated as:

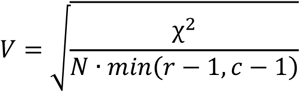

Where χ^2^ is the Pearson chi-squared statistic derived from a contingency table cross-tabulating the region of interest assignments of the two solutions. The term 𝑁 represents the total number of observations (i.e., 292 ROIs). The terms 𝑟 and 𝑐 represent the total number of distinct clusters in the first and second partition, respectively (corresponding to the number of rows and columns in the contingency table). The resulting 𝑉 statistic ranges from 0 to 1, where 1 indicates perfect agreement between the two networks and 0 indicates complete independence. The Mean Cramér’s V for a specific partition *k* was calculated by averaging its pairwise 𝑉 scores against all other solutions (𝑘′ ≠ 𝑘), with higher values reflecting a more stable cluster solution.

#### 2.6.6 Mean percent agreement

Mean Percent Agreement was calculated to evaluate the stability of each cluster solution. A greedy bipartite matching algorithm was used to map corresponding clusters between solutions based on maximal spatial overlap.

To compare a target cluster partition against another solution, a contingency table was generated cross-tabulating the region of interest assignments. The matching algorithm iteratively identified the cluster pair with the highest absolute number of shared ROIs until all possible clusters were paired. For any two cluster solutions, the Percent Agreement (𝑃𝐴) was calculated as:

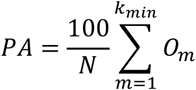

where 𝑁 represents the total number of ROIs in the parcellation scheme, and 𝑂_m_ denotes the number of overlapping ROIs within the 𝑚-th optimally matched cluster pair. The summation occurs over 𝑘_min_, representing the maximum number of possible matched pairs between the two solutions. To measure the overall robustness of a specific solution, its spatial boundaries were evaluated against all other solutions for *k*. For example, the *k=3* solution was iteratively compared against *k=2*, *k=4*, up to *k=15*. From these pairwise calculations, the Mean Percent Agreement was calculated, with higher values reflecting a cluster solution that remains stable across varying levels of granularity.

### 2.7 Examining cluster evolution across time points

Consensus clustering was applied independently at each timepoint to identify stable, group-wise cluster solutions for each age. This creates a challenge for the examination of functional network evolution across development however, as for a given *k* (e.g., *k = 5*), the precise ROI makeup (ROI identity/location; number of ROIs) of each cluster will differ across timepoints. We addressed this by calculating the voxel-weighted Jaccard index between all clusters, as a metric of spatial similarity, then applying the Kuhn-Munkres (Hungarian) algorithm; a matching algorithm the finds the optimal global assignment (i.e. identify the global optimal cluster matches between timepoints). Using the volume of an ROIs (voxel-weighted) in the calculation of the Jaccard index ensures that spatial similarity is not disproportionately skewed by the presence or absence of small anatomical regions. The impact of using equally weighted ROIs **(i)** versus ROIs weighted by number of voxels **(ii)** on metrics is shown in **Figure 3**.

**Figure 3.**
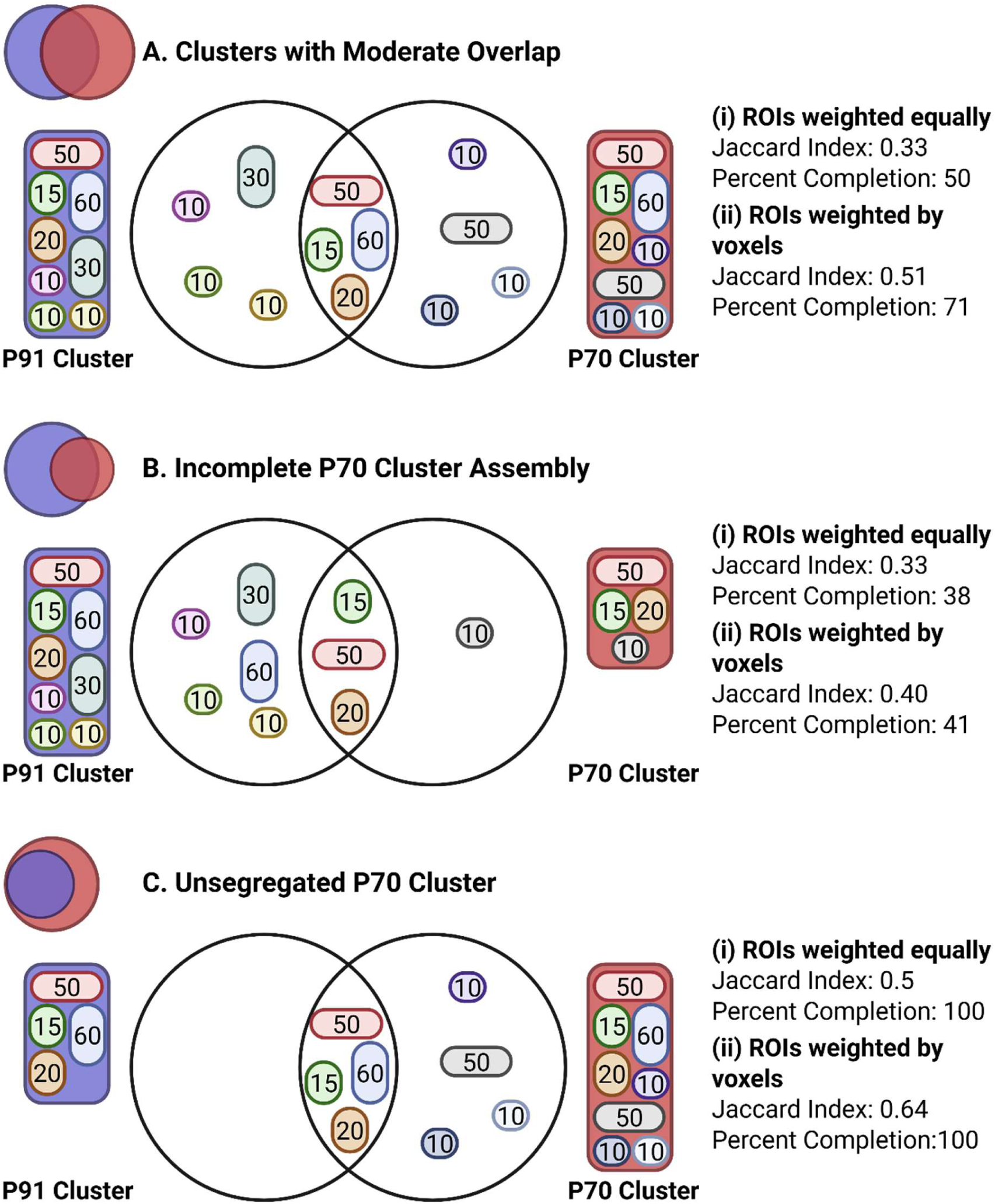
Examples of Jaccard index and percent completion and the impact of voxel weighting. The left cluster (purple) represents a mature cluster (P91) and the right cluster represents an immature cluster at an earlier age (e.g. P70). Shapes represent ROIs, with numbers inside indicating the number of voxels in that ROI. **(A)** Shows two clusters of similar size with a moderate overlap. **(B)** Shows incomplete assembly where the immature cluster (P70) is smaller than the mature cluster (P91), as many ROIs have not joined this cluster yet. **(C)** Shows an unsegregated immature cluster (P70), which contains all the ROIs in the mature (P91) cluster, but contains many others that are not present in the mature cluster. For each example, a comparison of Jaccard index and percent completion using ROIs that are **(i)** equally weighted or **(ii)** weighted by the number of voxels in each ROI.

#### 2.7.1 Spatial similarity metric: voxel-weighted Jaccard index

As the 292 anatomical ROIs vary in physical volume, spatial overlap between adjacent ages was measured using voxel-weighted Jaccard index (𝐽_w_). For a cluster 𝐶_t1_ at age 1 and a cluster 𝐶_t2_ at age 2, 𝐽_w_ is defined as:

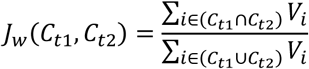

where 𝑉_i_ represents the total number of voxels comprising region 𝑖. The numerator calculates the total volume of the ROIs shared between two clusters (i.e. the intersection), while the denominator calculates the total voxel volume of all unique ROIs present in either cluster (i.e. the union).

This calculation results in a value between 0 and 1. A higher index value indicates the physical boundaries of the cluster (functional network) are similar at both ages, while a low 𝐽_w_ value indicates volumetric mismatch. In a developmental context, low spatial overlap may reflect one of two states: either the juvenile network has not yet recruited anatomical volumes (incomplete assembly, decreasing the intersection; **Figure 3B**), or regions are still contained within a larger, unsegregated immature cluster, which inflates the union between the clusters (**Figure 3C**).

#### 2.7.2 Identifying matching clusters across age using the Kuhn-Munkres algorithm

To find the optimal cluster match between ages (scanning timepoints), the Kuhn-Munkres (Hungarian) algorithm was used. First, the 𝐽_w_ similarities were converted into a dissimilarity/cost matrix for each adjacent timepoint pairing. Each element 𝐷(𝐶_t1_, 𝐶_t2_) within the matrix was defined as 1 − 𝐽_w_(𝐶_t1,_, 𝐶_t2_).

The Kuhn-Munkres (Hungarian) algorithm was then applied to this dissimilarity matrix at each developmental step. By evaluating all possible cluster permutations, this algorithm identifies the global combination of cluster pairs that minimises the total matrix cost. This stepwise approach was carried out in reverse chronological order: P91 – P70, P70 – P49, P49 – P35 and P35 – P28. This ensured the optimal identification of precursor functional networks across development. Cluster matching steps are summarised in **Figure 4**.

**Figure 4.**
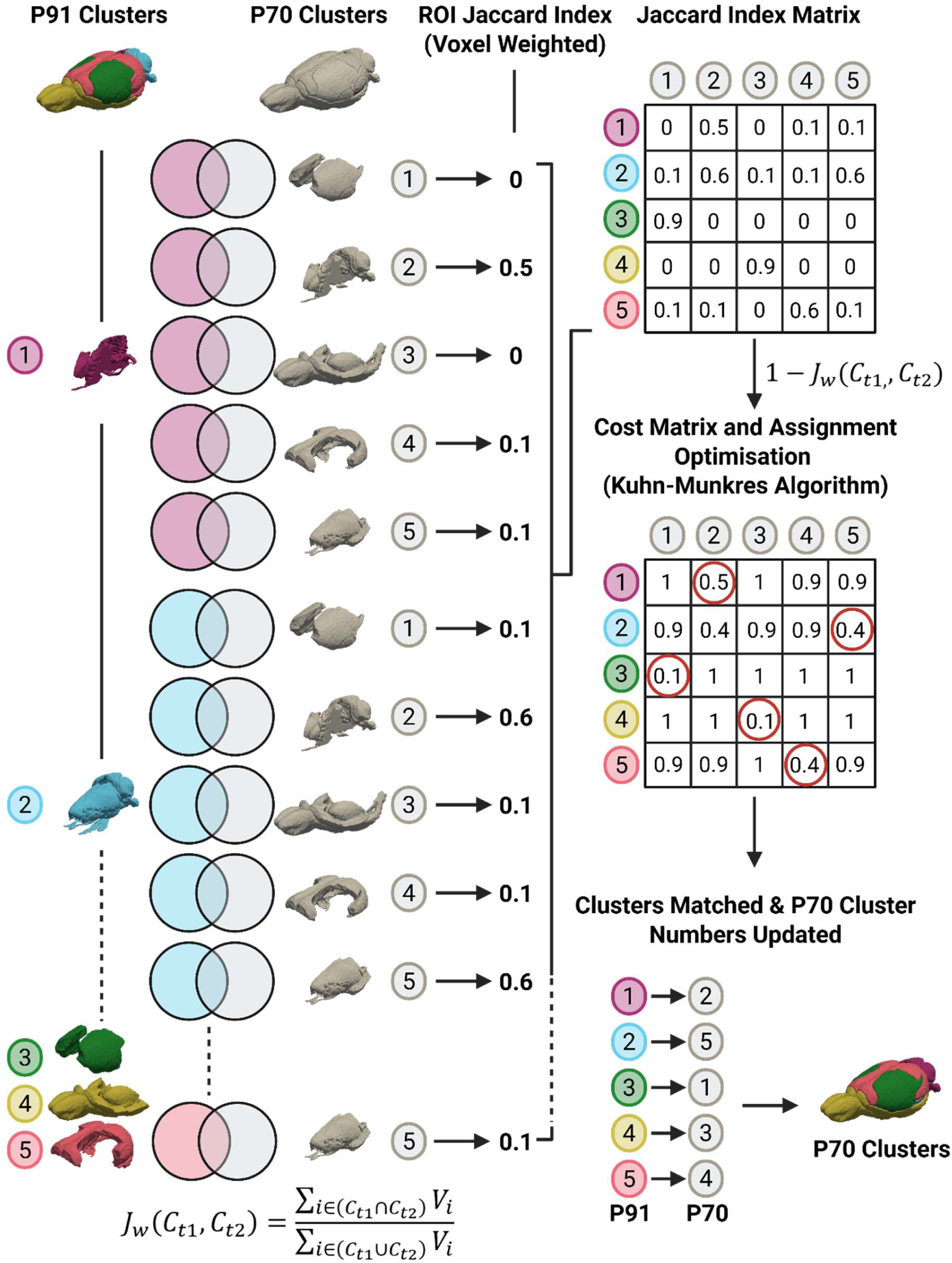
Summary of the steps used to identify precursor clusters over multiple timepoints. In order to track how clusters mature over time, the voxel-weighted Jaccard index between P91 and P70 was calculated for all clusters for a given *k* (k = 5 is shown). The weighted Jaccard index matrix between each cluster was then converted to a cost/dissimilarity matrix by subtracting values from 1. The Kuhn-Munkres/Hungarian algorithm was then used to find the optimal global cluster matches between ages. The cluster labels were then updated for P70 based on the optimal global matches to P91. Using the updated labels for the P70 clusters, these steps were repeated for P70 to P49, P49 to P35, and P35 to P28, using the updated labels between each step.

**Figure 5.**
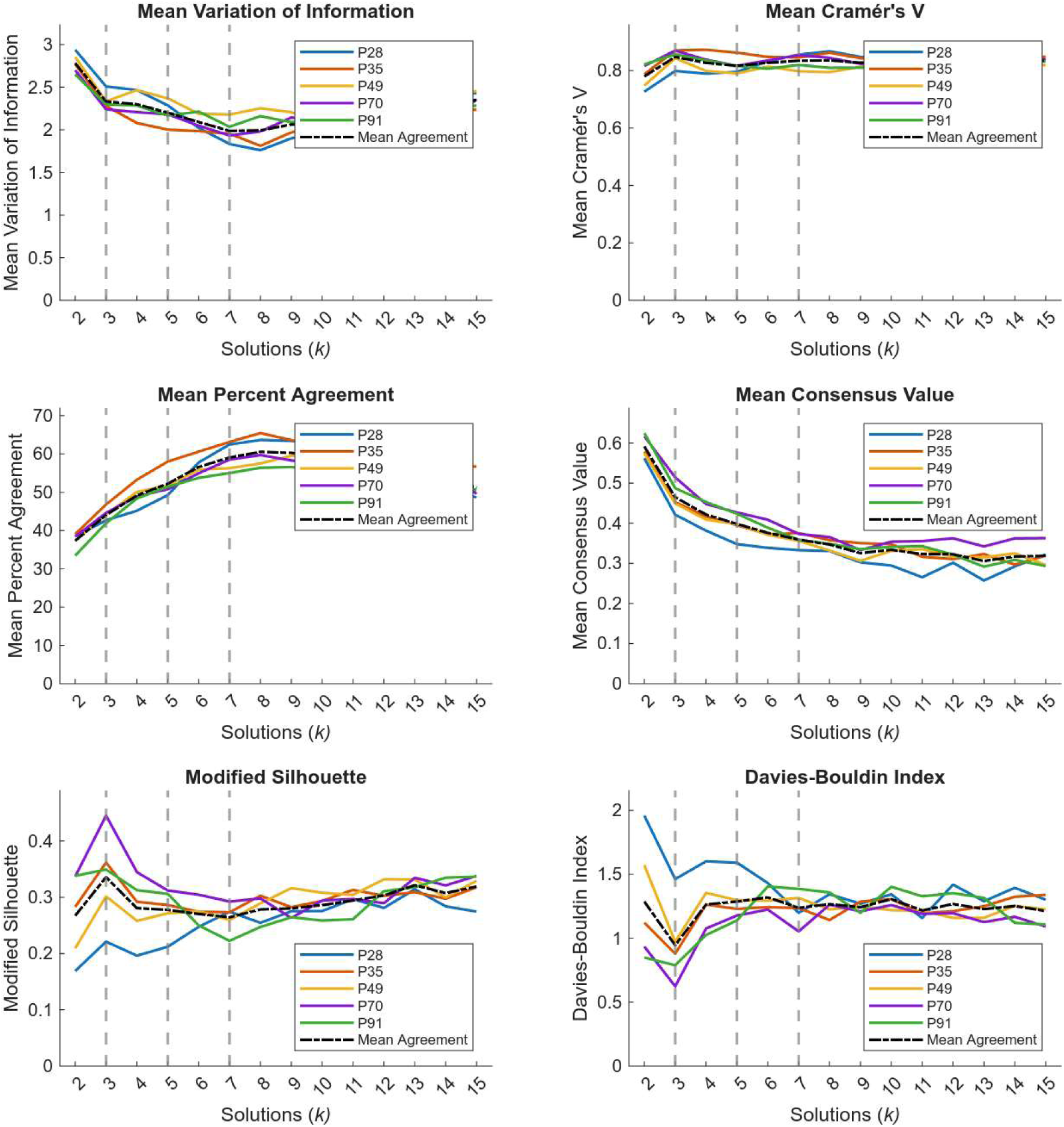
Cluster quality metrics. To identify clusters for further analysis, a series of cluster quality metrics were calculated for *k*=2 to *k*=15. Metrics include mean variation of information mean Cramer’s V, mean percent agreement, modified silhouette, Davies-Bouldin index and mean consensus. Solid lines represent each timepoint, with the broken line representing the mean agreement across timepoints.

### 2.8 Quantifying developmental trajectories

Once similar clusters had been identified across timepoints by the above approach, we used the last timepoint (P91) as a reference to investigate maturation. The voxel-weighted Jaccard index was calculated between P91 and earlier timepoints (P28, P35, P49 and P70; **Table 1**). Voxel-weighted percent completion was also calculated between P91 and earlier timepoints. Cluster colours for brain renderings (**Figure 6**), organisation of the connectivity matrices (**Figure 8**), node colours in the spring embeddings (**Figure 9**), and networks used in the calculation of system segregation and participation coefficient (**Figure 10**) are all determined by cluster membership at P91.

**Figure 6.**
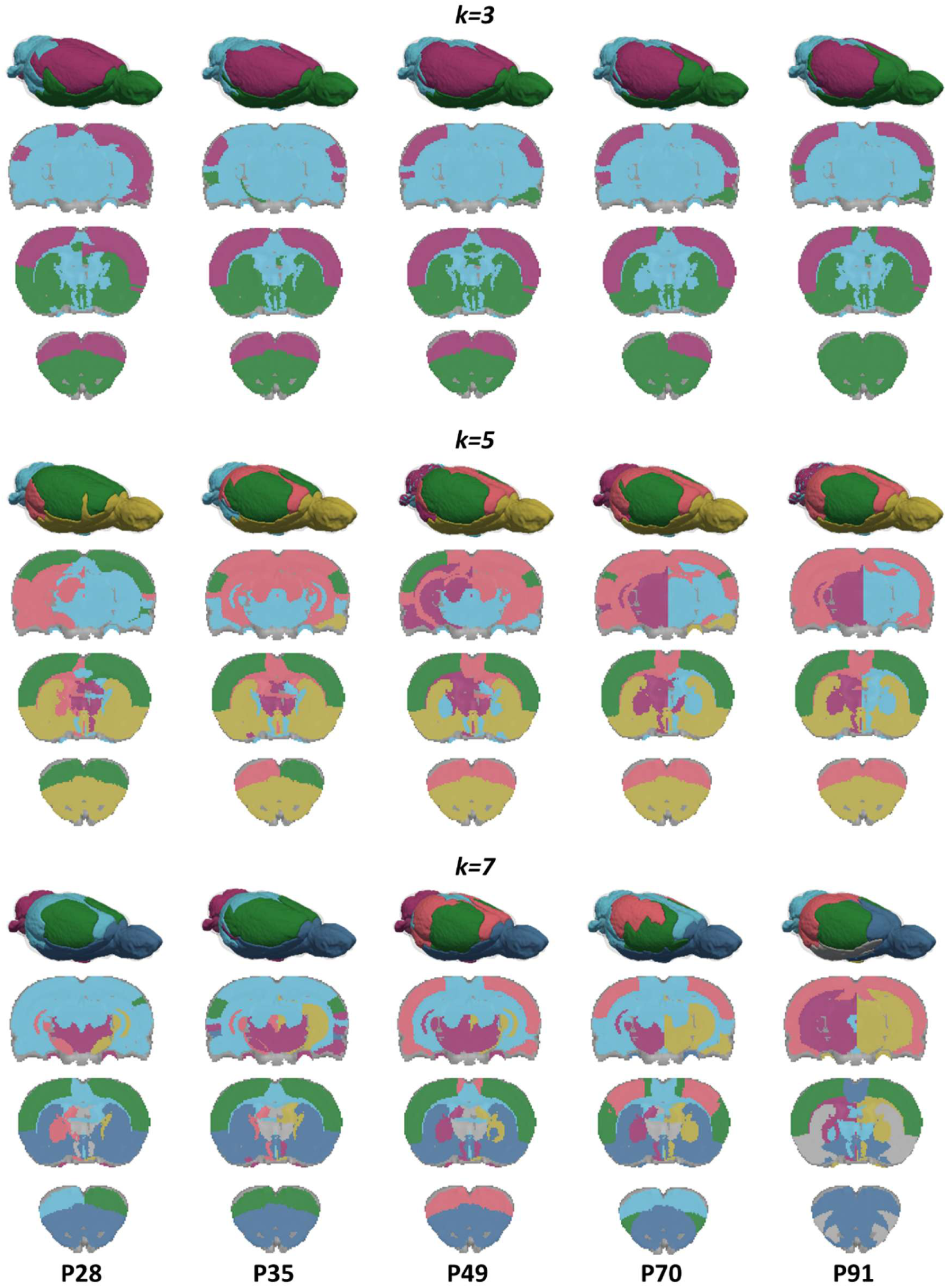
Brain mapping of clusters across development. Maps of functional networks from the juvenile period (P28) to adulthood (P91) are visualised across three solutions of *k* (*k=3, 5,* and *7*). To track the developmental lineage and spatial reorganisation of these networks, all ROIs are colour-coded based on their mature adult assignments at P91. Three-dimensional surface renderings of the brain networks were created using ITK-SNAP and ParaView^47^. Two-dimensional axial slices were generated using Nilearn at coordinates Z = −5, 0, and 5 mm. Colour schemes for each cluster are consistent across all figures and tables.

**Figure 7.**
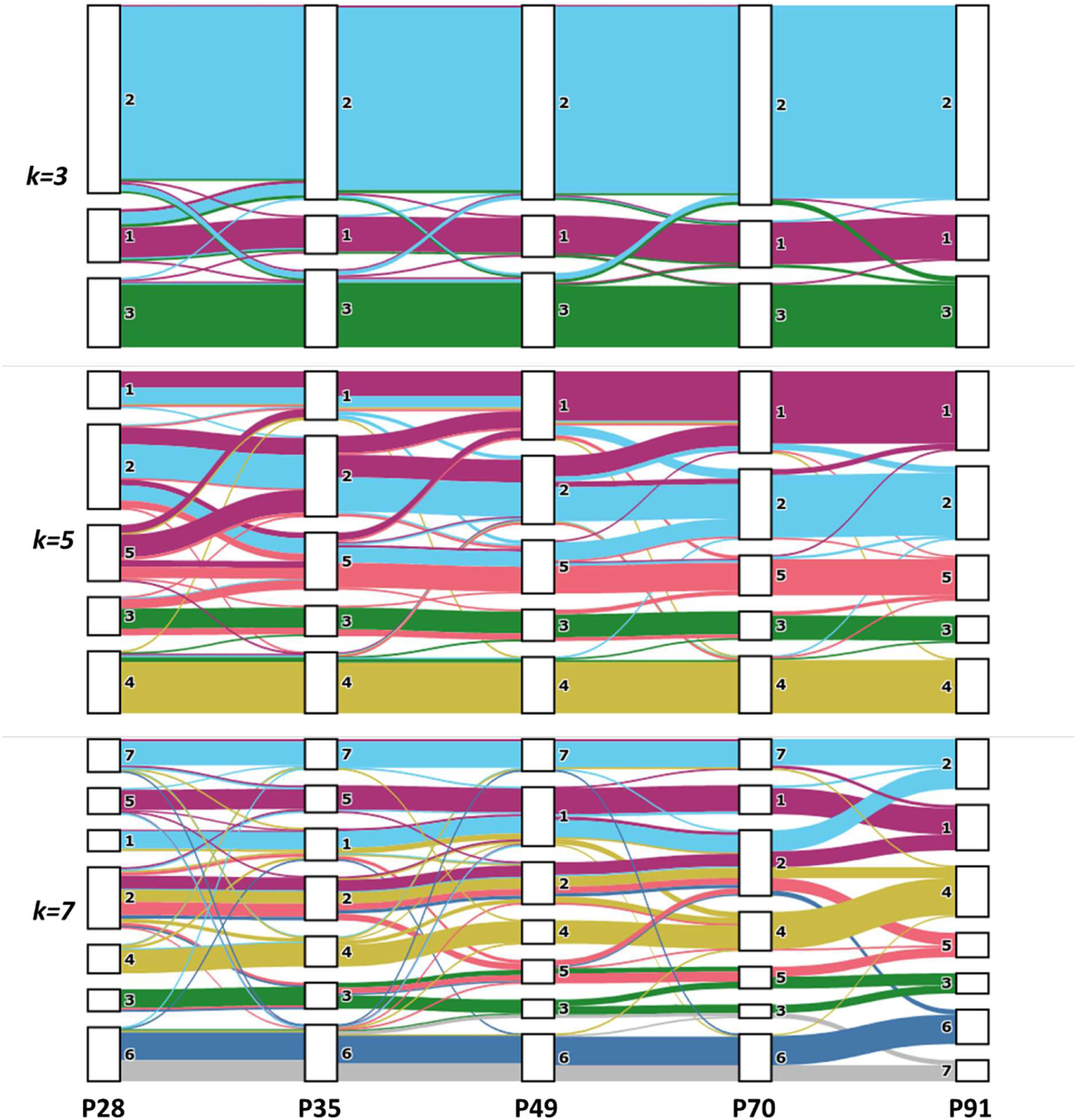
Longitudinal lineage tracking and developing functional connectome for *k=3, 5,* and *7.*. Sankey diagrams illustrating how ROIs change cluster membership over time. To map the developmental lineage of mature networks, a stepwise backward-matching analysis was adopted. As summarised in Figure 4, empirically derived clusters at each age were matched to the adjacent younger timepoint (e.g., P91 matched to P70, P70 to P49) using a voxel-weighted Jaccard index dissimilarity matrix, and optimised via the Kuhn-Munkres (Hungarian) algorithm.

**Figure 8.**
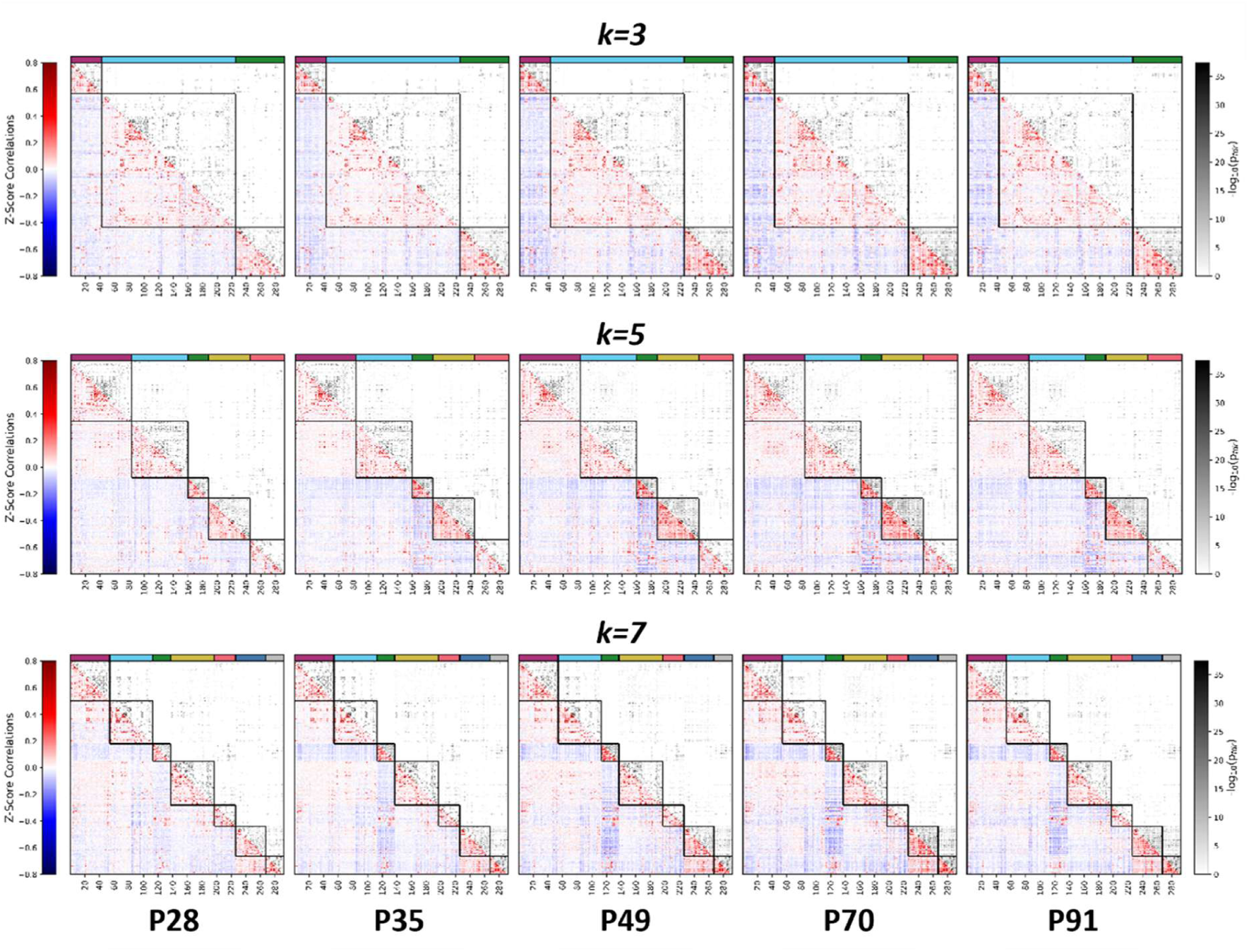
Within-network and between-network connectivity across development. Mean functional connectivity (FC) matrices and corresponding statistical significance maps show the connectome from P28 to P91. Matrices were ordered to match the P91 cluster assignments for each solution of *k*. The colour bars positioned along the top indicate the specific mature cluster identity of each ROI. The bottom triangle displays the group-average Fisher Z-transformed correlation values (red = positive connectivity, blue = negative connectivity). The top triangle displays the corresponding statistical significance of the connections at each age. For every edge, a one-sample t-test was performed across subjects to identify connections significantly greater than zero. P-values were corrected for multiple comparisons using the Benjamini-Hochberg FDR correction. Values are presented as −log_1O_(p_fdr_) where darker colours indicate greater statistical significance.

**Figure 9.**
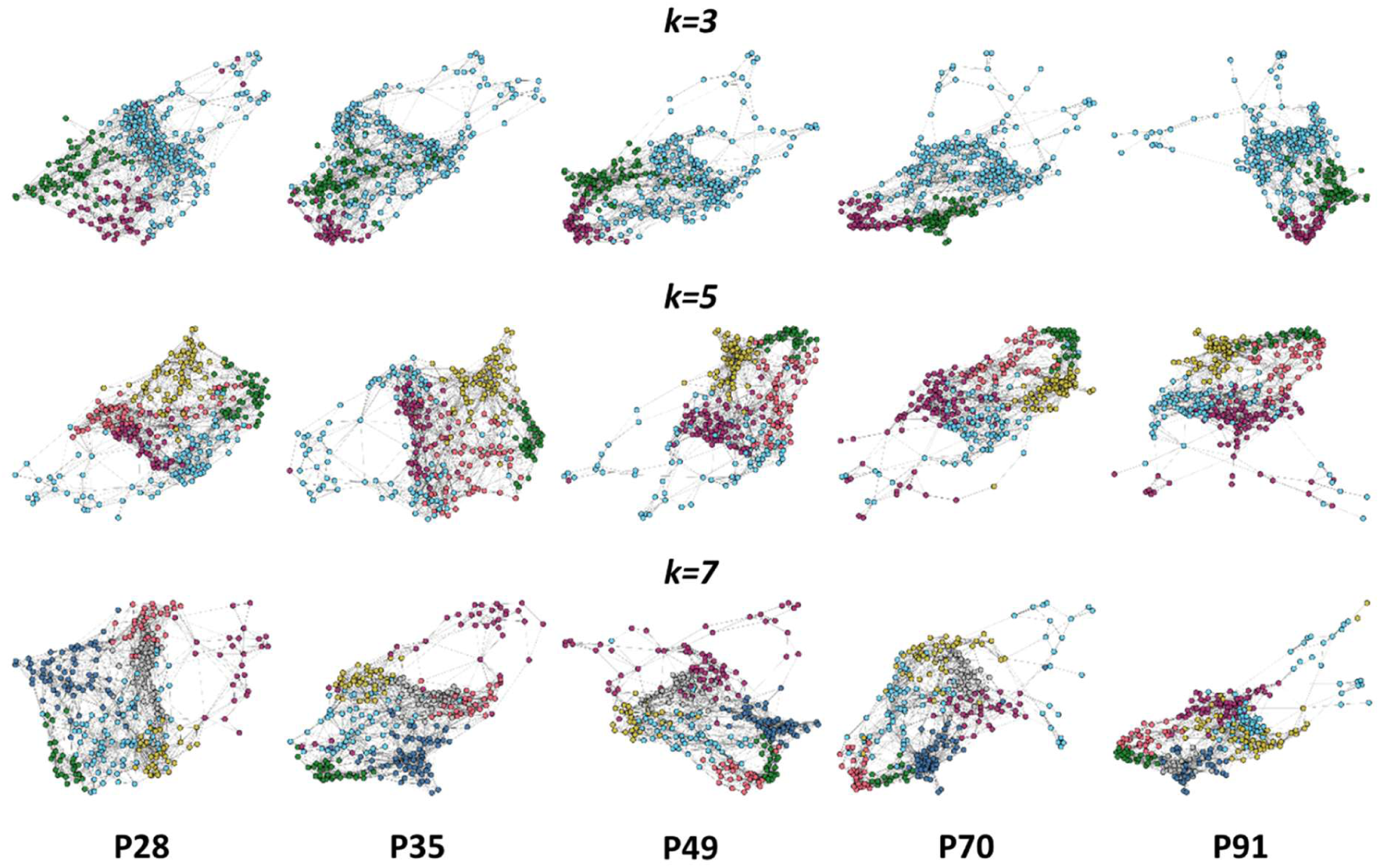
Topological reorganisation of the functional connectome across adolescence. Images highlight changes in functional brain networks from the juvenile period (P28) to adulthood (P91) using a force-directed spring layout algorithm. To generate a sparse matrix for embedding, the mean Fisher Z-transformed correlation matrices for each age group were thresholded to retain only the strongest 5% of functional connections (95th percentile). Nodes are coloured based on their final mature network assignment at P91 for *k=3*, *k=5* and *k=7*.

**Figure 10.**
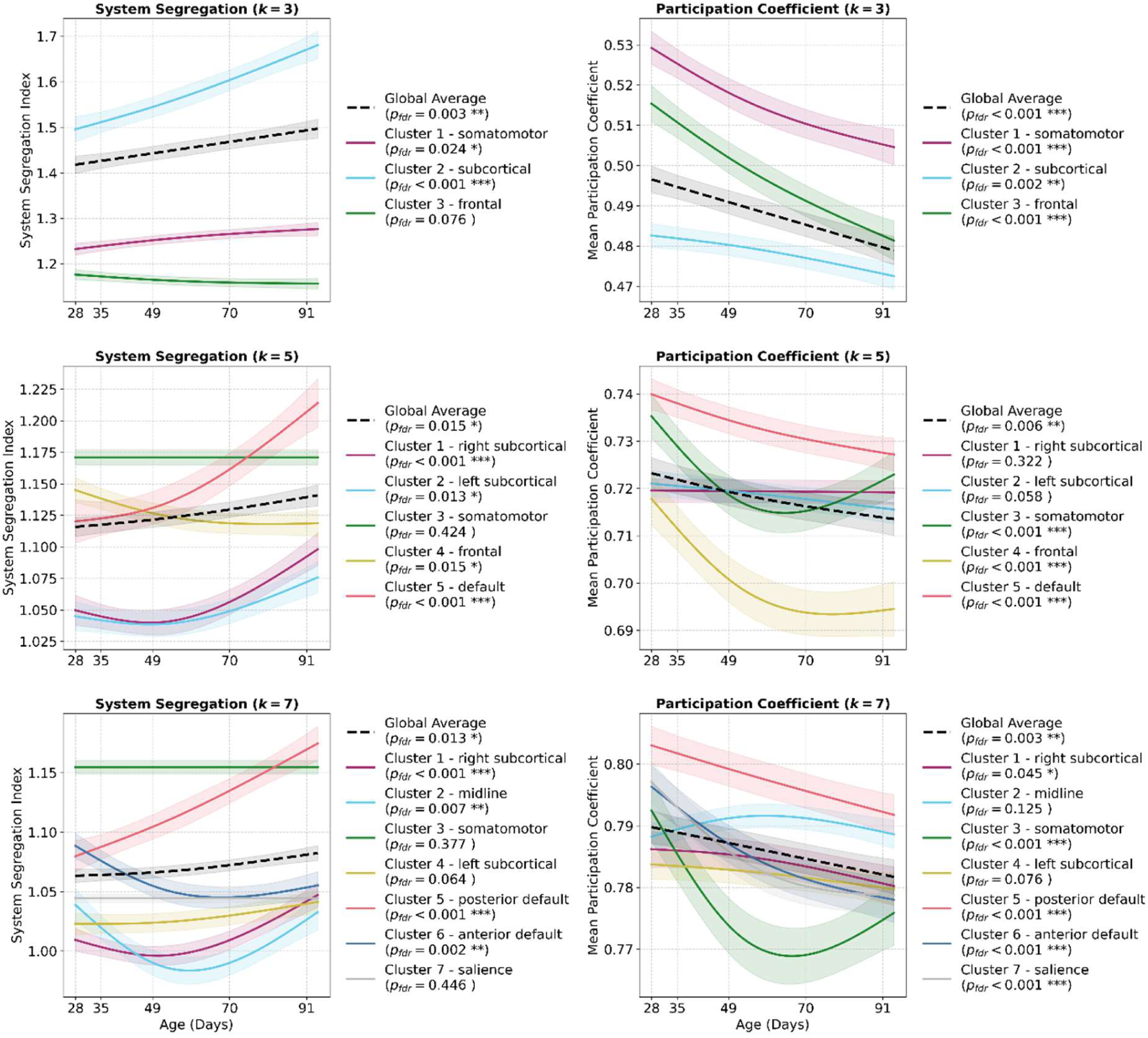
Longitudinal trajectories of network integration and segregation across adolescence. Developmental trajectories of global and cluster-specific topological metrics from juvenile (P28) to adulthood (P91), evaluated across three solutions for *k* (*k=3*, *k=5* and *k=7*). System segregation (left column) quantifies the difference between mean within-network and between-network connectivity. Calculations were performed on each subject’s fully signed, unthresholded Fisher Z-transformed connectivity matrices to incorporate both positive and negative edge weights^39^. Participation coefficient (right column) is a measure of network integration reflecting how uniformly a node’s connections are distributed across all clusters. To account for negative correlations, the participation coefficient was calculated separately for positive and negative edge weights, and the mean of these two values was used to represent the overall integration score for each respective network^39^. For both metrics, age-related developmental changes were modelled using a generalised additive mixed models to capture both linear and non-linear trajectories, with subject included as a random effect. The black dashed line represents the global average across the entire brain (mean of all clusters for that solution of *k*), while coloured lines depict the specific trajectories of individual functional clusters (coloured to match their mature P91 assignments). Shaded bands denote the 95% confidence intervals. and all *p*-values were corrected for multiple comparisons using the Benjamini-Hochberg FDR correction across the 36 independent tests. Asterisks (*) adjacent to the cluster labels denote trajectories that showed a statistically significant main effect of age, where p_fdr_<0.05*, p_fdr_<0.01** and p_fdr_<0.001***.

**Table 1.** Weighted Jaccard index and percent completion across development relative to P91 for *k=3*, *K=5* and *K=7*. Weighted Jaccard index (Left) using the clusters at P91 as a reference. The Jaccard index is defined as the size of the intersection divided by the size of the union of the sample sets. For a given cluster, each timepoint was compared to the P91 cluster; a higher index indicates greater similarity between sample sets. Weighted percent completion (Right) of each cluster using P91 as a reference. This describes how complete each cluster is. Both metrics were weighted by the number of voxels in each ROI to prevent smaller regions from disproportionately affecting cluster similarity. The weighted mean in the table accounts for the number of ROIs in each cluster. R: right; L: left; A: anterior; P: posterior.

| Weighted Jaccard Index |  |  |  |  | Weighted Percent Completion |  |  |  |  |
| --- | --- | --- | --- | --- | --- | --- | --- | --- | --- |
| k=3 |  |  |  |  |  |  |  |  |  |
| Cluster ID | P28 | P35 | P49 | P70 | Cluster ID | P28 | P35 | P49 | P70 |
| 1<br>Somatomotor | 0.544 | 0.683 | 0.775 | 0.875 | 1<br>Somatomotor | 80.9 | 78.65 | 88.24 | 96.26 |
| 2<br>Subcortical | 0.837 | 0.901 | 0.932 | 0.959 | 2<br>Subcortical | 87.68 | 96.98 | 97.7 | 99.51 |
| 3<br>Frontal | 0.797 | 0.788 | 0.81 | 0.874 | 3<br>Frontal | 85.61 | 86 | 86.41 | 87.76 |
| Mean | 0.726 | 0.791 | 0.839 | 0.903 | Mean | 84.73 | 87.21 | 90.78 | 94.51 |
| Weighted Mean | 0.768 | 0.829 | 0.869 | 0.92 | Weighted Mean | 85.8 | 90.51 | 92.92 | 95.84 |
| k=5 |  |  |  |  |  |  |  |  |  |
| Cluster ID | P28 | P35 | P49 | P70 | Cluster ID | P28 | P35 | P49 | P70 |
| 1<br>R-Subcortical | 0.06 | 0.097 | 0.48 | 0.645 | 1<br>R-Subcortical | 6.82 | 10.61 | 60.74 | 95.34 |
| 2<br>L-Subcortical | 0.54 | 0.437 | 0.507 | 0.623 | 2<br>L-Subcortical | 81.54 | 66.24 | 63.99 | 65.1 |
| 3<br>Somatomotor | 0.467 | 0.741 | 0.759 | 0.921 | 3<br>Somatomotor | 87.91 | 92.09 | 93.94 | 97.3 |
| 4<br>Frontal | 0.881 | 0.896 | 0.924 | 0.939 | 4<br>Frontal | 97.58 | 97.34 | 97.29 | 97.78 |
| 5<br>Default | 0.298 | 0.529 | 0.483 | 0.91 | 5<br>Default | 39.12 | 78.81 | 67.64 | 93.44 |
| Mean | 0.449 | 0.54 | 0.631 | 0.808 | Mean | 62.59 | 69.02 | 76.72 | 89.79 |
| Weighted Mean | 0.464 | 0.537 | 0.621 | 0.799 | Weighted Mean | 63.32 | 69.35 | 75.49 | 88.12 |
| k=7 |  |  |  |  |  |  |  |  |  |
| Cluster ID | P28 | P35 | P49 | P70 | Cluster ID | P28 | P35 | P49 | P70 |
| 1<br>R-Subcortical | 0.09 | 0.093 | 0.24 | 0.457 | 1<br>R-Subcortical | 18.01 | 19.79 | 51.04 | 47.46 |
| 2<br>Midline | 0.013 | 0.001 | 0.004 | 0.332 | 2<br>Midline | 3.88 | 0.31 | 0.86 | 84.35 |
| 3<br>Somatomotor | 0.842 | 0.713 | 0.85 | 0.501 | 3<br>Somatomotor | 95.91 | 95.91 | 90.47 | 54.98 |
| 4<br>L-Subcortical | 0.233 | 0.349 | 0.264 | 0.596 | 4<br>L-Subcortical | 23.42 | 34.98 | 26.38 | 60.34 |
| 5<br>P-Default | 0 | 0 | 0.469 | 0.291 | 5<br>P-Default | 0 | 0 | 63.15 | 44 |
| 6<br>A-Default | 0.415 | 0.407 | 0.444 | 0.441 | 6<br>A-Default | 73.46 | 73.42 | 73.46 | 73.75 |
| 7<br>Salience | 0 | 0 | 0 | 0 | 7<br>Salience | 0 | 0 | 0 | 0 |
| Mean | 0.228 | 0.223 | 0.324 | 0.374 | Mean | 30.67 | 32.06 | 43.62 | 52.13 |
| Weighted Mean | 0.231 | 0.233 | 0.319 | 0.398 | Weighted Mean | 31.32 | 33.1 | 43.19 | 55.14 |

#### 2.8.1 Voxel-weighted Jaccard index compared to P91

To measure the overall spatial overlap of clusters at each developmental timepoint, the voxel-weighted Jaccard index (𝐽_w_) was evaluated against the mature P91 reference set. While an identical formula from the step-wise matching analysis was used, each juvenile timepoint (P28, P35, P49, P70) was directly compared to P91 rather than its nearest age-neighbour. Unlike percent completion, the Jaccard index heavily penalises the inclusion of extraneous regions, as these unpruned ROIs inflate the union denominator. Therefore, it is possible for a juvenile cluster to reach 100% completion relative to its adult counterpart while still exhibiting a low Jaccard Index due to the presence of additional/extraneous ROIs not present in the mature network.

#### 2.8.2 Voxel-weighted percent completion compared to P91

To quantify the physical assembly of functional networks across development, we calculated a voxel-weighted percentage completion metric for each juvenile cluster relative to its mature P91 reference state. This metric measures the total volumetric “functional footprint” recruited at each developmental timepoint compared to the final timepoint. Scaling the assembly to each network’s adult volume normalises the developmental trajectories across all clusters, allowing for direct comparison of maturation rates between spatially disparate systems. Furthermore, tracking these completion trajectories (0–100%) facilitates the identification of critical neurodevelopmental windows during which functional networks experience increased volumetric expansion.

For a juvenile cluster 𝐶_t_ at timepoint 𝑡, the Percent Completion relative to the mature reference cluster 𝐶_P91_ is calculated as:

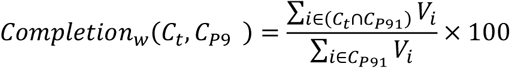

where 𝑉_i_ denotes the voxel count of ROI 𝑖. This formula divides the anatomical volume (in voxels) of the regions correctly assigned to the cluster at the current developmental age by the total target anatomical volume of the mature adult network at P91.

#### 2.8.3 Stepwise ROI membership change calculation

To identify specific age transitions characterised by the greatest degree of functional reorganisation, we calculated the step-wise ROI membership change. Following the alignment of cluster labels via the Kuhn-Munkres (Hungarian) algorithm, this metric quantifies the absolute proportion of ROIs that change their network membership between adjacent timepoints 𝑡_1_ and 𝑡_2_:

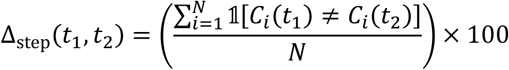

where 𝑁 represents the total number of ROIs in the parcellation scheme (292), and 𝟙 is an indicator function that equals 1 if the cluster assignment 𝐶_i_for region 𝑖 differs between timepoint 𝑡_1_and 𝑡_2_, and 0 if the assignment remains stable.

### 2.9 Visualising topology: Spring embedding

To visualise the topological architecture of the functional connectome across development, network embeddings were generated using a force-directed spring layout algorithm. Spring embedding is a commonly used method to display graph network data, such as brain networks^37^. In spring embedding algorithms, functional connections between nodes are treated as springs with a spring constant added which is relative to the connection strength between a pair of nodes. In our case, nodes were randomly placed on a 2D plane and the repellent force set to 0.9 (α) using *sknetwork* (0.33.3) in *python 3.10*. As connections between nodes are “spring loaded” they exert a force on each other. The algorithm then goes through 1000 iterations, adjusting the position of each node at every iteration to reduce the total energy of the system to the lowest possible energy state.

Importantly, spring embedding in naïve to the spatial location of nodes as it relies complete on correlation strength. Therefore, it provides information about the network in purely topological space. The mean Fisher Z-transformed matrices was thresholded to keep only the top 5% of connections (95^th^ percentile) to generate a sparse matrix. The largest connected component was then used as the input for spring embedding. This was repeated for each timepoint. Nodes were coloured based on the mature P91 cluster set for each *k* solution (3, 5, 7).

### 2.10 Quantifying developmental change: System segregation

System segregation measures the difference in mean within-system connectivity and mean between-system connectivity as a proportion of within-system connectivity. In this case, system segregation for a given cluster was calculated using the mean connectivity within a cluster and mean connectivity between that cluster and every other cluster for that solution. The average of these was used to calculate the global system segregation for a cluster solution. As in Chan et al.^38^, system segregation was defined as:

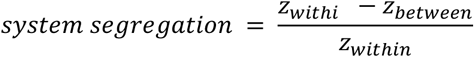

where 𝑧_within_ is the mean edge weight between nodes within the same cluster, and 𝑧_between_is the mean edge weight between nodes of one cluster to all nodes in other cluster (Adapted from *MATLAB* code at: https://github.com/mychan24/system_matrix_tools). Unthresholded Fisher Z-transformed matrices were used to calculate system segregation as in Tooley et al.^39^

### 2.11 Quantifying developmental change: Network integration

The participation coefficient is a measure of network integration that quantifies the diversity of the edges of a node across clusters and is sensitive to changes in network segregation in early life and across the lifespan^38–41^. For a given node 𝑖 participation coefficient 𝑃_i_ is defined as:

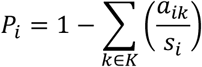

Where 𝑘 is a system in a set 𝐾 of systems (in this case defined as the mature P91 cluster solutions).

𝑎_ik_ is the positive (or negative) edge weights between node 𝑖 and nodes in system 𝑘, and 𝑠_i_ is the positive (or negative) strength of node 𝑖. The participation coefficient was calculated separately on negative and positive weights^39,42^.

The mean positive and negative participation coefficient for each participant’s network at each level of *k* was averaged to obtain a global measure of network integration for each *k*^39^.

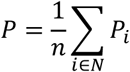

Where 𝑛 is the total number of nodes within network 𝑁.

### 2.12 Statistical assessment of developmental change: Generalised additive mixed model

To statistically assess the developmental effects on system segregation and participation coefficient (integration), we fitted a generalised additive mixed model (GAMM), which examines both linear and non-linear relationships between variables and age. This model included age as a smooth term, with subject ID as a random effect. Cubic splines were used as the basis for the smooth term as they provide computation efficiency and are suitable when data is sparse at the boundaries. Restricted maximum likelihood (REML) for the selection of smoothing parameters was used, which allows unbiased estimation of variance components in random effects models^43^. Models were run in *R* with code adapted from Li et al.^44^. Full model outputs are available in the **supplemental material 2**.

### 2.13 Statistical analysis

Generalised additive models (**Figure 10**) were run in *R* version 4.1.2, using packages; *dplyr* (1.0.8) and *mgcv* (1.8.39). To control for multiple testing across all comparisons for the graph metrics (36 comparisons), the Benjamini-Hochberg FDR correction was applied in python 3.10 using *statsmodels* (0.14.5). Statistical significance was defined as 𝑝_fdr_< 0.05.

A one-sample *t-test* was performed on every edge in the mean Fisher Z-transformed matrix for each age (**Figure 8**) to identify connections significantly greater than zero in *python* 3.10, using *statsmodels* 0.14.5. P-values were corrected for multiple comparisons using the Benjamini-Hochberg FDR correction in *statsmodels* 0.14.5. Values are presented as −log_1O_(𝑝_fdr_).

## 3. Results

### 3.1 Optimal cluster identification

To capture the organisation of brain networks at different levels, we took a multi-resolution approach, focusing on cluster solutions *k=3*, *k=5*, and *k=7*. The selection of these solutions was informed by the cluster quality metrics (**Figure 5**). The coarse cluster solution of *k=3* was supported by internal validity metrics; specifically, a low Davies-Bouldin index was observed across nearly all age groups, indicating compact and well-separated clusters. This is further supported by a peak in the modified silhouette, suggesting strong cluster cohesion, and a high mean consensus value, reflecting robust cross-subject stability. As an intermediate solution, *k*=5 was selected as it falls on the ’elbow’ of the mean consensus curve. Investigation of *k*=5 is further supported by the Cramer’s V, which remains stable at this level. Finally, *k*=7 was identified as the optimal fine-grained network organisation. The mean variation of information reaches a trough, indicating relatively homogeneous and distinct clusters. Moreover, the Davies-Bouldin index stabilises at this point.

### 3.2 The mature networks

To understand what each cluster represents in terms of functional systems, we first investigated the mature adult solutions (P91), as these networks serve as the lens through which we view developmental changes^45^.

#### 3.2.1 Coarse partition

A cluster solution of 3 (*k = 3*) provided a macroscale view of the brain architecture, offering information on broad network systems. In the mature brain, Cluster 1 (purple; **Figure 6**) consisted of 42 ROIs and was comprised of left and right somatosensory and motor areas. Regions included the primary and secondary motor cortices, primary and secondary somatosensory cortices, primary and secondary visual cortices auditory cortices, parietal association areas, and insular cortices. Cluster 2 (light blue; **Figure 6**) was the largest cluster and consisted of 183 left and right subcortical and limbic ROIs. Regions included the medial cortical regions: cingulate area, retrosplenial area; the hippocampal formation (CA1, CA2, CA3, dentate gyrus, subiculum and entorhinal cortices), the majority of the thalamic nuclei; the hypothalamus; and hindbrain and midbrain regions (cerebellum, brainstem, pontine nuclei, and substantia nigra). Cluster 3 (green; **Figure 6**) was made up of 67 ROIs from frontal and subcortical brain areas. Prominent members included most of the prefrontal cortex (prelimbic, infralimbic, and orbital cortices); the striatum (caudate-putamen, nucleus accumbens, ventral striatum, ventral pallidum, globus pallidus); the limbic regions (amygdaloid area, claustrum, septal regions, basal forebrain); and the olfactory system (olfactory bulb, piriform cortex, endopiriform nucleus, lateral olfactory tract).

#### 3.2.2. Intermediate partition

The greater detail provided by *k=5* allowed the lateralisation of brain networks to be captured (Figure X.). Cluster 1 (purple; **Figure 6**) was comprised of 83 ROIs spanning primarily right subcortical structures. Cluster 2 (light blue; **Figure 6**) consisted of 77 ROIs, and nearly perfectly mirrored Cluster 1, with primarily left subcortical structures. Cluster 3 (green; **Figure 6**) was a bilateral somatomotor network made up of 28 ROIs; regions included the primary and secondary motor cortices, primary and secondary somatosensory cortices, granular, dysgranular, and agranular insular cortices. Cluster 4 (yellow; **Figure 4**) was a large bilateral frontal cluster consisting of 57 ROIs. It can be broadly separated into three sub-areas: the prefrontal cortex (prelimbic, infralimbic, and all orbital cortices); the striatum (caudate-putamen, nucleus accumbens, ventral pallidum, ventral striatum); and limbic and olfactory areas (amygdaloid area, olfactory bulb, piriform cortex, claustrum, endopiriform nucleus, and basal forebrain). Cluster 5 (red; **Figure 6**) consisted of 47 ROIs located in the medial and posterior cortex, with some subcortical structures, and was consistent with the rodent default network^46^. This cluster can be broadly separated into the hippocampal formation, medial cortical areas (cingulate and retrosplenial cortices), and posterior sensory areas, namely the primary and secondary visual and auditory cortices and temporal association area.

#### 3.2.3. Fine partition

This mature P91 cluster solution for *k=7* displayed clear bilateral symmetry and distinct cortical-subcortical boundaries. Much like *k*=5, a solution of *k*=7 split subcortical areas into distinct left and right clusters. At this solution, Cluster 1 (purple; **Figure 6**) contained 54 right subcortical ROIs, while Cluster 4 (yellow; **Figure 6**) contained 60 left subcortical ROIs. The main constituents of these clusters were the hippocampal formation, major white matter tracts (corpus callosum, fimbria, stria terminalis), extensive thalamic nuclei (anteroventral, ventromedial, laterodorsal), and midbrain/basal structures (substantia nigra, globus pallidus, ventral tegmental area). Cluster 2 (light blue; **Figure 6**) was a midline cluster of 59 ROIs consisting of the hypothalamus, periaqueductal grey, septal region, cerebellum, pons, medulla (inferior olive), and midline thalamic nuclei. Cluster 3 (green; **Figure 6**) is a small bilateral cluster with 24 ROIs and contained somatomotor regions such as the primary and secondary motor cortices, primary and secondary somatosensory cortices, parietal association areas, granular insula. Cluster 5 (red; **Figure 6**) is a posterior cortical network made up of 29 ROIs, with its major regions being the primary and secondary visual and auditory cortices, along with the retrosplenial cortex, entorhinal cortex (medial and lateral), perirhinal and postrhinal cortices. This cluster could be considered a posterior unit of the rodent default network, combined with posterior sensory regions. Cluster 6 (dark blue; **Figure 6**) was a bilateral frontal cluster consisting of 41 ROIs and featured frontal default network regions. Major regions included the prelimbic area, infralimbic area, anterior cingulate, orbital cortices (ventral, lateral, medial), nucleus accumbens, ventral pallidum, and olfactory bulb. Cluster 7 (grey; **Figure 6**) was a frontal lateral cluster of 25 ROIs, broadly resembling the rodent salience network, with major nodes including the caudate-putamen, amygdala, piriform cortex (primary olfactory), claustrum and insula. For each value of k, comprehensive ROI lists are included in the **supplemental material 2**.

### 3.3. Assembly of the adult network over time

At the macro-scale (*k=3*), the tripartite architecture of the brain is largely established by P28, with 85.8% of all ROIs (voxel-weighted mean percent completion) matching the mature adult organisation (weighted Jaccard index of 0.743 relative to P91). Over development, weighted percent completion and weighted Jaccard index gradually increased, reaching 95.84% and 0.92 respectively, by P70.

However, longitudinal tracking revealed a period of functional boundary refinement across adolescence. Most notably, hubs of the rodent default network; the retrosplenial and cingulate areas decouple from the somatomotor network to join the mature frontal network (Cluster 3) at P35 and P70 respectively. Similarly, the Insular areas segregate from prefrontal-limbic circuits to join the dedicated somatomotor network (Cluster 1) between P35 and P70, demonstrating delayed functional specialisation of the cortex. In line with this, the greatest number of ROIs changing cluster membership occurred between P28 and P35 (11.3%; 33/292; **Table 1**), and this stabilised to 4.79% (14/292), 6.16% (18/292) and 3.77% (11/292) for transitions between subsequent timepoints.

Sequentially tracking *k=5* using longitudinal analysis revealed that trajectories of functional network maturation are heterogeneous. The frontal network (Cluster 4) is almost completely established by P28, with a stable weighted percent completion of 97.29% to 97.78% across timepoints and weighted Jaccard indices between 0.881 and 0.939 (**Table 1**). Similarly, the somatomotor network (Cluster 3) contains almost all the regions of the adult network by P28 with a weighted percent agreement of 87.91% that rises marginally to 97.30% by P70. In contrast, networks associated with higher-order functions, like the default network (Cluster 5) exhibited significant maturational change across the developmental window of this study. At P28, Cluster 5 is particularly immature, exhibiting only 39.12% completion and a weighted Jaccard index of 0.298. It then expands its functional volume throughout adolescence by recruiting hippocampal, entorhinal, and visuospatial areas that were previously functionally tethered to the somatomotor network in the juvenile brain. The greatest number of ROIs changed membership at the P28 to P35 transition (31.16%; 91/292; **Table 2**). A second major transition was observed between P49 and P70 (26.03%; 76/292; **Table 2**), primarily driven by lateralisation of the left and right subcortical regions into their own distinct clusters (Clusters 1 and 2).

**Table 2.** Number of ROIs that change cluster between ages. The total number of ROIs clustered was 292.

| Stepwise ROI Membership Change |  |  | 647 |
| --- | --- | --- | --- |
| *k=3* |  |  | 648 |
| Transition | ROIs Changed | % of Total ROIs |  |
| P28 - P35 | 33 | 11.3 |  |
| P35 - P49 | 14 | 4.79 | 650 |
| P49 - P70 | 18 | 6.16 |  |
| P70 - P91 | 11 | 3.77 |  |
| *k=5* |  |  |  |
| Transition | ROIs Changed | % of Total ROIs |  |
| P28 - P35 | 91 | 31.16 |  |
| P35 - P49 | 57 | 19.52 |  |
| P49 - P70 | 76 | 26.03 |  |
| P70 - P91 | 30 | 10.27 |  |
| *k=7* |  |  |  |
| Transition | ROIs Changed | % of Total ROIs |  |
| P28 - P35 | 44 | 15.07 |  |
| P35 - P49 | 91 | 31.16 |  |
| P49 - P70 | 69 | 23.63 |  |
| P70 - P91 | 135 | 46.23 |  |

Tracking of the more fine-grained *k*=7 parcellation revealed that brain maturation at this functional scale is more complex, and is characterised by network fractionation during adolescence. While global Jaccard index and percent completion increase with age, as observed for *k*=3 and *k*=5, stepwise changes in cluster membership were greatest at the transition from P70 to P91, with 46.23% of ROIs (135/292; **Table 2**) changing membership. This was primarily driven by the late-stage emergence of the salience network and lateralisation of the subcortical network. The salience network (Cluster 7) does not exist as an independent entity at earlier timepoints; rather, its constituent ROIs are completely subsumed within the frontal network (Cluster 6) until late adolescence. It is only in the final developmental window (P70-P91) that regions including the amygdaloid area, piriform cortex and insular cortices functionally uncouple from the prefrontal network (Cluster 6) to form Cluster 7. Across development, stepwise volatility is heavily driven by the asymmetric lateralisation of the deep subcortical/brainstem cores (Clusters 1 and 4). Longitudinal tracking showed that the left-hemisphere subcortical core (Cluster 4) established a strictly lateralised spatial boundary as early as P28. In contrast, the right-hemisphere subcortical core (Cluster 1) operates as a functionally fused, bilateral hub throughout early development. It is not until P70 that the Cluster 1 right-hemisphere subcortical core undergoes a large-scale spatial reorganisation, losing its left-hemispheric and cerebellar nodes, which accounts for a significant portion of the network volatility observed in the mid-adolescent window. Both networks then undergo a final wave of region recruitment at P91 to finalise their mature, mirrored thalamocortical boundaries. The flow of ROIs between clusters at each age are shown in **Figure 7**.

### 3.4 Changes in within-network and between-network connectivity over development

Visual examination of the mean connectivity matrices across development (**Figure 8**) suggests a pattern of increasing within-network connectivity, coupled with decreasing between-network connectivity. These observations are reflected in the force-directed spring embeddings of the functional networks (*k*=3, 5, and 7; **Figure 9**). As spatial proximity in these layouts is driven exclusively by functional connection strength, the shape of the graph physically illustrates the developmental strengthening and weakening of connections. At the first two developmental timepoints (P28 and P35), the connectome is presented as a large, continuous, and globally integrated topology. Nodes belonging to the same future adult network (indicated by node colour) are distributed loosely throughout the structure, reflecting a diffuse developmental state with low modularity. This indicates that internal connections within specific functional networks were not yet strong enough to overcome the dense web of weak, between-network connections that physically bind the immature brain into diffuse, weakly differentiated networks. During adolescent timepoints (P49 and P70), the network architecture undergoes spatial consolidation. Driven by the targeted weakening of between-network connections and the simultaneous strengthening of within-network connections, resulting in nodes belonging to the same functional cluster moving closer together. By adulthood (P91), the connectome is much more modular. The large, diffuse juvenile web has transitioned into more dense, isolated networks, with nodes of the same cluster closer together.

### 3.5 Developmental trajectories of network topology

The spring embedding analysis allows for visualisation and qualitative analysis of the evolution of topological reorganisation across development. We also quantified these observable patterns using metrics that quantify network segregation (system segregation) and integration (participation coefficient), and tested for the significance of linear and non-linear age effects using the generalised additive mixed model (GAMM). **Figure 10** shows the longitudinal trajectories of these metrics at the global average and cluster-specific level, across the three solutions for *k* (*k=3, k=5 and k=7*).

#### 3.5.1 Macro-scale pruning and system maturation

At the global network level, system segregation displayed an upward trajectory from P28 to P91 across all spatial scales (*k=3: edf =1.16, F=4.9, p_fdr_ =0.003; k=5: edf=1.06, F=3.02, p_fdr_ =0.015, and k=7: edf=1.19, F=3.49, p_fdr_ =0.013*). In line with the results from system segregation, participation coefficient, a measure of integration, underwent an age related decrease across all solutions for *k* explored (*k=3: edf=1.42, F=13.74, p_fdr_ <0.001; k=5: edf=1.16, F=4.35, p_fdr_ =0.006, and k=7: edf=1.16, F=5.22, p_fdr_ =0.003*). These changes in within- and between-network connectivity are consistent with the observations made for the connectivity matrices and spring embeddings.

At the coarsest spatial scale, (*k*=3), the brain exhibited a significant decrease in integration over adolescence, with all three clusters displaying a lower participation coefficient between P28 to P91. System segregation remained stable in the prefrontal-limbic network (*Cluster 3: edf=1.03, F=1.60 p_fdr_ =0.066*) but showed significant age-related increases in the somatomotor (*Cluster 1; edf=1.06, F=2.59, p_fdr_ =0.024*) and subcortical (*Cluster 2; edf=1.45 F=13.88, p_fdr_ <0.001*) networks. The macroscale view highlights that development is marked by highly integrated juvenile brain networks that become more refined (less diffuse; more segregated) over time.

#### 3.5.2 Late-stage assembly of the default network

At higher spatial resolutions, development changes in higher order brain networks, such as the rodent default network, became apparent. At *k*=5, the rodent default network (Cluster 5) displayed a strong, significant increase in segregation that accelerated notably after P49 (*edf=1.45, F=8.03, p_fdr_=<0.001*), paired with a steady, significant decrease in integration over time (*edf=1.33, F=7.65; p_fdr_ <0.001*).

At the highest cluster solution, *k*=7, the default network is split between two clusters, with Cluster 5 representing posterior components such as the retrosplenial cortex, while Cluster 6 represents the anterior subunit of the network and included hallmark regions of the rodent default network, namely the cingulate cortices and prelimbic cortices. Interestingly, we see subunit specific development of the default network, with the posterior default network (Cluster 5) demonstrating a steep, stable increase in segregation (*edf=1.47, F=15.01; p_fdr_ =<0.001*) and a significant decrease in integration over time (*edf= 1.25, F=7.29, p_fdr_ <0.001*). In contrast, the frontal components of the default network (Cluster 6) exhibited a significant decrease in segregation at early timepoints (*edf=1.71, F=6.74; p_fdr_ =0.002*), alongside a sharp initial drop in integration that begins to level off by adulthood (*edf=1.59, F=15.45; p_fdr_ <0.001*). The divergence between these two subunits suggests that while the posterior default network steadily consolidates over development, the anterior default network undergoes changes in early adolescence before stabilising into its mature adult configuration.

#### 3.5.3 Non-linear maturation of the somatomotor network

The somatomotor network exhibited a distinctly different trajectory than the default network. The somatomotor network (Cluster 3, for both *k=5* and *k=7*) displayed stable system segregation across the developmental window captured by this study, with no significant changes observed from P28 to P91 (*k=5: edf=0.0005, F=0.002, p_fdr_ =0.424; k=7: edf=0.0013, F=0.001, p_fdr_ =0.377*). This lack of change in segregation over time suggests that the functional boundaries of the primary motor and sensory cortices are already mature at the juvenile stage (P28). Interestingly, however, the integration of this network follows a significant U-shaped curve, with the participation dropping sharply at early timepoints before reversing course and significantly increasing at P70 and P91 *(k=5: edf=1.85, F=12.23, p_fdr_ <0.001; k=7: edf=1.86 F=15.61, p_fdr_ <0.001*).

#### 3.5.4 Delayed segregation of the lateralised subcortical networks

Finally, longitudinal tracking of the subcortical networks revealed a delayed but structurally symmetric maturation process. At *k=5,* the subcortical regions are divided into distinct left and right hemispheric clusters (Clusters 1 and 2) from P70 onwards. Both of these bilateral clusters exhibited largely stable integration across development (*Cluster 1: edf=0.11, F=0.06, p_fdr_ =0.322; Cluster 2: edf=0.86, F=1.63 p_fdr_ =0.058*), but demonstrated significant, late-emerging increases in segregation that only became evident after P49, which is in agreement with the emergence of these clusters in the lineage-tracking analysis (*Cluster 1: edf=1.70 F=8.13, p_fdr_ <0.001; Cluster 2:edf=1.53, F=4.18 p_fdr_ =0.013*). This pattern is almost perfectly replicated at *k=7*, where the left (Cluster 1) and right (Cluster 4) subcortical clusters emerged at P70. At *k=7*, the left subcortical core (Cluster 1) showed an increase in segregation after P49 (*edf=1.78 F=9.42 p_fdr_ <0.001*), paired with a significant drop in integration (*edf 1.04 F=2,16 p_fdr_ =0.045*), while its right hemisphere counterpart (Cluster 4) maintained a similar but statistically non-significant trajectory (*edf=0.85 F=1.41 p_fdr_ =0.076*). This late-stage shift suggests a critical window during mid-to-late adolescence for the lateralisation of subcortical brain networks.

## 4. Discussion

In this study, we aimed to map the longitudinal spatiotemporal maturation of functional brain networks from the juvenile, pre-pubertal period (P28) to early adulthood (P91) in the Wistar rat. By analysing the developmental evolution of functional networks across several scales (*k=3, k=5 and k=7),* we revealed that the rodent brain shows differentiated, network-specific developmental changes. Our analyses showed that functional network development was characterised by a transition from a more diffusely connected juvenile state to a more segregated, refined, and functionally specialised adult state. Network-specific trajectories replicated the hierarchical sensory-association axis observed in humans, with sensory networks maturing first and association networks maturing last ^38,39,48^. Our goal was to identify a robust, sex-balanced functional network architecture in the Wistar rat that can enable further studies that leverage the translational power of resting state fMRI. To accelerate further translational work, our data (https://openneuro.org/datasets/ds007859), parcellations (https://codeberg.org/mclooned/rat_developmental_clustering/src/branch/main/network_parcellati ons), and code (https://codeberg.org/mclooned/rat_developmental_clustering) are openly available.

### 4.1 Translational homology

Using consensus clustering, we identified mature adult rat functional networks and how their precursors mature over time using a sex-balanced dataset. Across all solutions of *k* investigated, a somatomotor network was readily identifiable, which included visual and auditory components at *k*=3 (Cluster 1), but became strictly composed of sensory and motor regions at *k*=5 and *k*=7 (Custer 3). This network is frequently observed across mice and rats - it is commonly termed the lateral cortical network, and overlaps with the somatomotor network in humans ^49^. This network displayed a high level of developmental stability, with its functional boundaries and segregation largely established by P28. This is consistent with well-established cross-species evidence that primary sensory networks are phylogenetically prioritised to mature before higher order networks^7^.

Homologs of the default network were also identified at *k*=5 and *k*=7. At *k*=5, Cluster 5 resembles the default network and can be broadly separated into the hippocampal formation, medial cortical areas including the cingulate area and retrosplenial cortex, and posterior visual and auditory areas. This is in broad agreement with other studies that have described the rodent default network homolog^46,49^. Interestingly, when the number of brain clusters is increased to 7 (*k*=7), we see the mature default network splitting into a frontal and posterior subnetworks. While anterior and posterior subcomponents of the default network have been identified in humans^50^, it has been reported that the rodent default network shows greater activity in frontal regions when compared with the human default network, which shows more balanced activity between posterior and frontal regions ^49^. The homolog of the default network was one of the last brain networks to mature, with the homolog at *k*=5 only reaching relative maturity by P70, reflecting the delayed maturation of higher order networks.

Interestingly, at *k*=7, we identified a homolog of the salience network, consisting of the insular cortices, amygdaloid area, and piriform cortex. This network only emerged at P91 after branching off from Cluster 6 (frontal default network subunit), suggesting it only displayed sufficient synchronous activity to be clustered together late in development. Of note, this network, as well as all other networks at *k*=7 display remarkable bilateral symmetry, with clear subcortical-cortical boundaries and the left and right sides mirroring each other near perfectly.

The use of longitudinal functional neuroimaging in rodents to investigate changes in brain structure and function over time has increased in recent years ^51–53^. However, to our knowledge, only one other longitudinal development study spanning juvenile to early adulthood has been carried out – in awake male Long Evans rats^54^. The authors noted that changes in functional connectivity were most pronounced between juvenile and late adolescence, suggesting this is a critical period for circuit development. Our findings are in agreement with this; development of higher order brain networks such as the default and salience network in our study was observed after P49. Similarly, we also observed lateralisation of subcortical brain networks between P49 and P70. Furthermore, Ma et al.^54^ noted that functional connectivity in somatomotor regions was largely established by the juvenile period – a pattern we also observed ^54^.

### 4.2 The topological and spatial trajectory of brain development

Changes in the topological reorganisation of the adolescent brain was captured by the inverse trajectories of the participation coefficient and system segregation. At early time points, the clusters presented as a highly integrated networks with high levels of cross talk between them. This was reflected in elevated participation coefficients across nearly all spatial scales. These functional networks underwent significant spatial consolidation as the animals progressed through adolescence, however, observed as significant age-related decreases in global integration, paired with increases in system segregation. These macroscale network changes likely reflect well-established changes taking place at the cellular level. Histological studies have shown that the juvenile brain displays a high density of synaptic connections that undergoes pruning throughout adolescence and into adulthood ^8,9^. As the brain matures, the active pruning of weak, unused, or irrelevant out-of-network connections, combined with the myelination of connections between intra-network hubs results in less physical connection between neural systems ^13,55^. This process drives functional shifts from a “local to distributed” connectome, resulting in highly segregated modular brain architecture^56,57^.

In line with these underlying processes, the percentage completion trajectories revealed a highly a heterogeneous assembly timeline. The large cortical and subcortical nodes of networks established their functional footprints early in life, whereas higher-order satellite regions such as the associative cortices were the last to functionally synchronise (i.e. be clustered together). This highlights that networks assemble progressively by recruiting distant anatomical nodes to form the mature adult boundaries.

While functional networks are a feature of the mammalian brain, it should be noted that humans have a richer brain topography than rodents. Mapping homologous brain networks from humans to non-human primates or rodents requires the presence of synchronous activity in phylogenetically conserved brain regions such as limbic and somatomotor cortical areas, or evolutionarily-ancient precursors of the medial prefrontal cortex like the anterior cingulate cortex^58^. Thus, higher order networks such as the default network are often only be partially recapitulated in non-human species^59–61^.

It is plausible that higher-order cognitive networks such as the dorsal-attention, temporoparietal or executive control networks are human-specific^62,63^. This is largely due to the fact that these networks are anchored to higher-order cortical areas that have undergone significant evolutionary expansion and are thus highly specialised in humans. In non-human primates, a candidate for the network involved in executive function has been observed, but a rodent homolog is yet to be identified, although possible candidates have been observed^64–66^. In the case of the human temporoparietal and dorsal attention networks, no non-human primate or rodent homologs have been identified.

Consequently, rodent and non-human primate precursors of human higher-order cognitive networks are yet to be firmly identified.

### 4.3 Limitations and future directions

A methodological strength of this study was the implementation of a sequential, voxel-weighted Jaccard matching pipeline, optimised using the Kuhn-Munkres matching algorithm. This approach robustly addressed the challenge of identifying precursor networks across development, resolving the “moving target” issue when applying clustering at multiple timepoints, particularly for intermediate *k* solutions (*k=5 to k=10*). This approach also controlled for atlas-specific spatial biases.

Furthermore, calculating topological metrics on fully signed, unthresholded matrices ensured our measures of segregation and integration account for both positive and negative correlations^39^.

However, it must be acknowledged that enforcing a strict one-to-one mapping across developmental timepoints represents a methodological compromise. In biological reality, the developing brain is highly dynamic; functional networks likely do not strictly evolve in isolated, one-to-one linear trajectories. While an approach that does not use this constraint might theoretically capture these complex states more comprehensively, it would render the longitudinal calculation of network metrics such as percent completion mathematically intractable. Therefore, enforcing a direct lineage strikes a balance between capturing the biological reality of network maturation and maintaining the statistical interpretability to quantify these changes. Furthermore, our approach of clustering can identify distinct non-overlapping networks that are highly synchronised at rest. In reality, and as in humans^67^, it is improbable the networks in rodents are completely distinct from each other and may show functional dynamics over time and in a task-dependent manner. For example, both the rodent default network and salience network incorporate the anterior cingulate area^68,69^.

Nevertheless, establishment of these multi-scale functional networks in male and female rats opens several avenues for future research. Using these consensus-derived functional networks, we can investigate changes in networks that might be relevant to disease. For example, by using these networks to establish the basis for a normative model, we can investigate how stress or environmental perturbations might alter the developmental trajectory of the rodent default network from pre-puberty to early adulthood.

## 5. Conclusion

In conclusion, this study establishes a sex-balanced multi-resolution functional network atlas for the developing rat brain. By leveraging a longitudinal design, we characterised the evolution of these functional networks across development. Across metrics, including cluster assembly and completion, force-directed spring embeddings, connectivity matrices, and graph metrics, we observed a strengthening of within-network connections and a weakening of between-network connections. As has been observed in humans, adolescence is a critical window of boundary refinement, but with a significant developmental gradient from sensorimotor to higher cognitive networks, such that higher-order networks, such as the default network, mature later in development. Since these functional parcellations are anchored to the Waxholm anatomical atlas and were captured under a consensus-standardised sedation and acquisition protocol, our functional architecture provides a generalisable structure and nomenclature for the preclinical imaging community, analogous to the Yeo 7-network parcellation in humans. This ultimately enables the investigation of functional network integrity in rodent models of disease and early life adversity, accelerating the discovery of translatable biomarkers for human neurological and psychiatric conditions.

## Data and Code availability

The imaging data used in this study are formatted according to the Brain Imaging Data Structure and are publicly available on OpenNeuro ds007859; https://openneuro.org/datasets/ds007859.

Clustering solutions in the population average space and SIGMA 2.0 rat brain space for each *k* and age are available at https://codeberg.org/mclooned/rat_developmental_clustering/src/branch/main/network_parcellati ons. The complete code to reproduce the analysis presented here are available at https://codeberg.org/mclooned/rat_developmental_clustering.

## AI statement

The authors declare that generative artificial intelligence has not been used for the writing of this manuscript, nor for the creation of figures, nor their corresponding legends.

