## supplemental material 1 for "Longitudinal consensus clustering reveals the functional architecture of the developing rat brain"

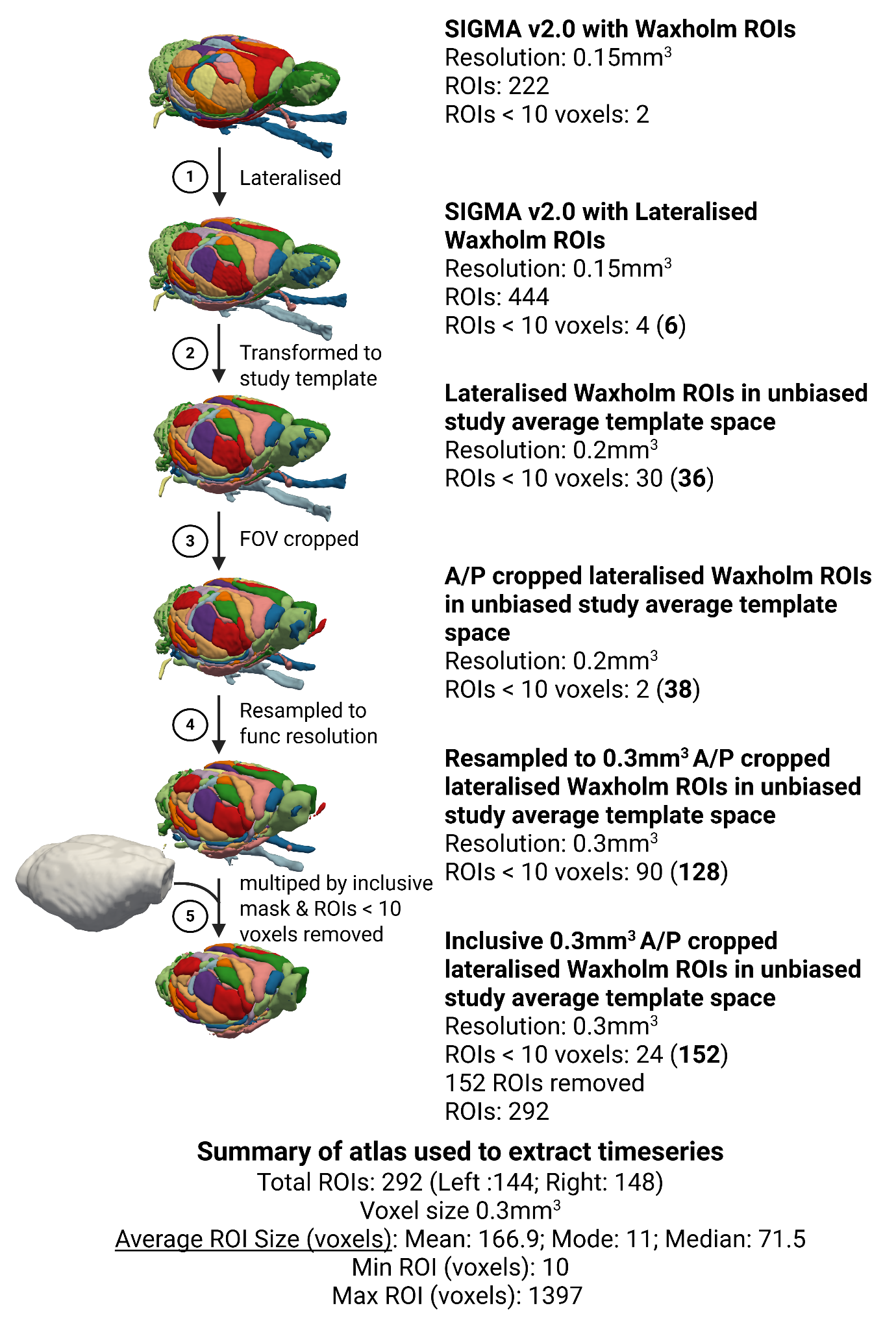
**Altas information**

| **ROIs in 0.3mm atlas population average** | | |
| --- | --- | --- |
| **ROI** | **NAME** | **VOXELS** |
| 1 | corticofugal_tract_and_corona_radiata_R | 504 |
| 4 | Molecular_cell_layer_of_the_cerebellum_R | 1397 |
| 5 | Cerebellum_unspecified_R | 1294 |
| 7 | inferior_cerebellar_peduncle_R | 43 |
| 10 | Cingulate_area_2_R | 135 |
| 32 | Entopeduncular_nucleus_R | 13 |
| 33 | Ventricular_system_unspecified_R | 395 |
| 34 | medial_lemniscus_unspecified_R | 20 |
| 36 | anterior_commissure_anterior_limb_R | 21 |
| 38 | ventral_hippocampal_commissure_R | 12 |
| 40 | Septal_region_R | 269 |
| 42 | optic_tract_and_optic_chiasm_R | 92 |
| 43 | Pineal_gland_R | 39 |
| 47 | Brainstem_unspecified_R | 1388 |
| 48 | Hypothalamic_region_unspecified_R | 259 |
| 50 | Superficial_gray_layer_of_the_superior_colliculus_R | 113 |
| 51 | Periaqueductal_gray_R | 277 |
| 52 | fornix_R | 25 |
| 55 | Deeper_layers_of_the_superior_colliculus_R | 395 |
| 56 | Periventricular_gray_R | 135 |
| 58 | Pontine_nuclei_R | 31 |
| 59 | fimbria_of_the_hippocampus_R | 205 |
| 61 | stria_medullaris_thalami_R | 24 |
| 62 | stria_terminalis_R | 43 |
| 64 | Glomerular_layer_of_the_accessory_olfactory_bulb_R | 15 |
| 65 | Glomerular_layer_of_the_olfactory_bulb_R | 85 |
| 66 | Olfactory_bulb_unspecified_R | 792 |
| 67 | corpus_callosum_and_associated_subcortical_white_matter_R | 1145 |
| 69 | inferior_colliculus_commissure_R | 24 |
| 71 | Interpeduncular_nucleus_R | 11 |
| 76 | spinal_trigeminal_tract_R | 164 |
| 77 | Frontal_association_cortex_R | 154 |
| 78 | middle_cerebellar_peduncle_R | 48 |
| 79 | transverse_fibers_of_the_pons_R | 27 |
| 82 | Basal_forebrain_region_unspecified_R | 758 |
| 93 | Bed_nucleus_of_the_stria_terminalis_R | 35 |
| 94 | Pretectal_region_R | 120 |
| 95 | Cornu_ammonis_3_R | 370 |
| 96 | Dentate_gyrus_R | 394 |
| 97 | Cornu_ammonis_2_R | 61 |
| 98 | Cornu_ammonis_1_R | 356 |
| 99 | Fasciola_cinereum_R | 38 |
| 100 | Subiculum_R | 201 |
| 108 | Postrhinal_cortex_R | 154 |
| 109 | Presubiculum_R | 162 |
| 110 | Parasubiculum_R | 125 |
| 112 | Perirhinal_area_35_R | 60 |
| 113 | Perirhinal_area_36_R | 121 |
| 114 | Medial_entorhinal_cortex_R | 339 |
| 115 | Lateral_entorhinal_cortex_R | 115 |
| 125 | 4th_ventricle_R | 113 |
| 128 | Dorsal_cochlear_nucleus_deep_core_R | 10 |
| 141 | lateral_lemniscus_unspecified_R | 55 |
| 142 | Inferior_colliculus_dorsal_cortex_R | 42 |
| 143 | Inferior_colliculus_central_nucleus_R | 107 |
| 145 | Inferior_colliculus_external_cortex_R | 208 |
| 146 | inferior_colliculus_brachium_R | 29 |
| 151 | Primary_auditory_area_R | 186 |
| 152 | Secondary_auditory_area_dorsal_part_R | 151 |
| 153 | Secondary_auditory_area_ventral_part_R | 145 |
| 157 | external_medullary_lamina_auditory_radiation_R | 14 |
| 180 | lateral_olfactory_tract_R | 65 |
| 181 | Piriform_cortex_layer_1_R | 217 |
| 182 | Piriform_cortex_layer_2_R | 53 |
| 183 | Piriform_cortex_layer_3_R | 226 |
| 184 | Nucleus_accumbens_core_R | 76 |
| 187 | Substantia_nigra_reticular_part_R | 99 |
| 188 | Substantia_nigra_compact_part_R | 10 |
| 192 | Nucleus_accumbens_shell_R | 91 |
| 193 | Ventral_pallidum_R | 71 |
| 195 | Globus_pallidus_external_medial_part_R | 20 |
| 196 | Ventral_tegmental_area_R | 38 |
| 197 | Caudate_putamen_R | 1020 |
| 198 | Globus_pallidus_external_lateral_part_R | 131 |
| 199 | Ventral_striatal_region_unspecified_R | 33 |
| 200 | Reticular_prethalamic_nucleus_unspecified_R | 99 |
| 204 | Pregeniculate_nucleus_R | 18 |
| 205 | Dorsal_lateral_geniculate_nucleus_R | 41 |
| 206 | Lateral_habenular_nucleus_R | 11 |
| 210 | Posterior_thalamic_nuclear_group_triangular_part_R | 12 |
| 211 | Parataenial_thalamic_nucleus_R | 13 |
| 214 | Anteroventral_thalamic_nucleus_dorsomedial_part_R | 12 |
| 215 | Anteroventral_thalamic_nucleus_ventrolateral_part_R | 26 |
| 219 | Reuniens_thalamic_nucleus_R | 31 |
| 221 | Ventromedial_thalamic_nucleus_R | 50 |
| 222 | Submedius_thalamic_nucleus_R | 10 |
| 227 | Ventral_posteromedial_thalamic_nucleus_R | 74 |
| 228 | Laterodorsal_thalamic_nucleus_dorsomedial_part_R | 19 |
| 229 | Laterodorsal_thalamic_nucleus_ventrolateral_part_R | 48 |
| 230 | Posterior_thalamic_nucleus_R | 68 |
| 231 | Ventrolateral_thalamic_nucleus_R | 68 |
| 232 | Mediodorsal_thalamic_nucleus_lateral_part_R | 15 |
| 233 | Mediodorsal_thalamic_nucleus_central_part_R | 11 |
| 235 | Zona_incerta_dorsal_part_R | 28 |
| 236 | Zona_incerta_ventral_part_R | 35 |
| 240 | Mediodorsal_thalamic_nucleus_medial_part_R | 22 |
| 242 | Paraventricular_thalamic_nuclei_anterior_and_posterior_R | 31 |
| 246 | Paracentral_thalamic_nucleus_R | 18 |
| 247 | Central_medial_thalamic_nucleus_R | 24 |
| 248 | Central_lateral_thalamic_nucleus_R | 11 |
| 249 | external_medullary_lamina_unspecified_R | 51 |
| 254 | Anteromedial_thalamic_nucleus_R | 33 |
| 267 | Parafascicular_thalamic_nucleus_R | 15 |
| 270 | superior_cerebellar_peduncle_and_prerubral_field_R | 16 |
| 281 | Subgeniculate_nucleus_R | 16 |
| 283 | Lateral_posterior_thalamic_nucleus_lateral_part_R | 18 |
| 285 | Lateral_posterior_thalamic_nucleus_mediorostral_part_R | 14 |
| 287 | Zona_incerta_caudal_part_R | 18 |
| 293 | Ventral_anterior_thalamic_nucleus_R | 15 |
| 294 | Ventral_posterolateral_thalamic_nucleus_R | 70 |
| 295 | Medial_geniculate_body_dorsal_division_R | 13 |
| 298 | Medial_geniculate_body_ventral_division_R | 40 |
| 400 | Ventrolateral_orbital_area_R | 72 |
| 401 | Lateral_orbital_area_R | 142 |
| 402 | Ventral_orbital_area_R | 83 |
| 403 | Medial_orbital_area_R | 44 |
| 404 | Dorsolateral_orbital_area_R | 114 |
| 405 | Prelimbic_area_R | 207 |
| 406 | Secondary_motor_area_R | 685 |
| 407 | Frontal_association_area_3_R | 150 |
| 408 | Primary_motor_area_R | 524 |
| 409 | Agranular_insular_cortex_ventral_area_R | 49 |
| 410 | Agranular_insular_cortex_dorsal_area_R | 90 |
| 411 | Cingulate_area_1_R | 194 |
| 412 | Claustrum_R | 73 |
| 413 | Infralimbic_area_R | 56 |
| 414 | Dysgranular_insular_cortex_R | 142 |
| 416 | Granular_insular_cortex_R | 136 |
| 417 | Primary_somatosensory_area_forelimb_representation_R | 335 |
| 418 | Primary_somatosensory_area_dysgranular_zone_R | 113 |
| 420 | Primary_somatosensory_area_face_representation_R | 559 |
| 422 | Secondary_somatosensory_area_R | 415 |
| 423 | Primary_somatosensory_area_hindlimb_representation_R | 165 |
| 424 | Agranular_insular_cortex_posterior_area_R | 88 |
| 425 | Primary_somatosensory_area_barrel_field_R | 547 |
| 427 | Retrosplenial_dysgranular_area_R | 256 |
| 429 | Primary_somatosensory_area_trunk_representation_R | 201 |
| 430 | Retrosplenial_granular_area_R | 279 |
| 432 | Parietal_association_cortex_lateral_area_R | 55 |
| 433 | Parietal_association_cortex_medial_area_R | 31 |
| 436 | Parietal_association_cortex_posterior_area_R | 58 |
| 442 | Primary_visual_area_R | 564 |
| 443 | Secondary_visual_area_lateral_part_R | 180 |
| 444 | Temporal_association_cortex_R | 147 |
| 448 | Secondary_visual_area_medial_part_R | 316 |
| 500 | Endopiriform_nucleus_R | 99 |
| 501 | Amygdaloid_area_unspecified_R | 411 |
| 502 | Nucleus_of_the_lateral_olfactory_tract_R | 24 |
| 1001 | corticofugal_tract_and_corona_radiata_L | 497 |
| 1004 | Molecular_cell_layer_of_the_cerebellum_L | 1249 |
| 1005 | Cerebellum_unspecified_L | 1185 |
| 1007 | inferior_cerebellar_peduncle_L | 42 |
| 1010 | Cingulate_area_2_L | 104 |
| 1032 | Entopeduncular_nucleus_L | 18 |
| 1033 | Ventricular_system_unspecified_L | 313 |
| 1034 | medial_lemniscus_unspecified_L | 33 |
| 1036 | anterior_commissure_anterior_limb_L | 27 |
| 1040 | Septal_region_L | 230 |
| 1042 | optic_tract_and_optic_chiasm_L | 85 |
| 1043 | Pineal_gland_L | 18 |
| 1047 | Brainstem_unspecified_L | 1339 |
| 1048 | Hypothalamic_region_unspecified_L | 284 |
| 1050 | Superficial_gray_layer_of_the_superior_colliculus_L | 99 |
| 1051 | Periaqueductal_gray_L | 220 |
| 1052 | fornix_L | 13 |
| 1055 | Deeper_layers_of_the_superior_colliculus_L | 394 |
| 1056 | Periventricular_gray_L | 110 |
| 1058 | Pontine_nuclei_L | 28 |
| 1059 | fimbria_of_the_hippocampus_L | 189 |
| 1061 | stria_medullaris_thalami_L | 26 |
| 1062 | stria_terminalis_L | 57 |
| 1064 | Glomerular_layer_of_the_accessory_olfactory_bulb_L | 17 |
| 1065 | Glomerular_layer_of_the_olfactory_bulb_L | 79 |
| 1066 | Olfactory_bulb_unspecified_L | 762 |
| 1067 | corpus_callosum_and_associated_subcortical_white_matter_L | 1197 |
| 1069 | inferior_colliculus_commissure_L | 11 |
| 1071 | Interpeduncular_nucleus_L | 15 |
| 1076 | spinal_trigeminal_tract_L | 189 |
| 1077 | Frontal_association_cortex_L | 138 |
| 1078 | middle_cerebellar_peduncle_L | 43 |
| 1079 | transverse_fibers_of_the_pons_L | 19 |
| 1082 | Basal_forebrain_region_unspecified_L | 749 |
| 1093 | Bed_nucleus_of_the_stria_terminalis_L | 46 |
| 1094 | Pretectal_region_L | 113 |
| 1095 | Cornu_ammonis_3_L | 370 |
| 1096 | Dentate_gyrus_L | 365 |
| 1097 | Cornu_ammonis_2_L | 56 |
| 1098 | Cornu_ammonis_1_L | 419 |
| 1099 | Fasciola_cinereum_L | 34 |
| 1100 | Subiculum_L | 202 |
| 1108 | Postrhinal_cortex_L | 162 |
| 1109 | Presubiculum_L | 146 |
| 1110 | Parasubiculum_L | 112 |
| 1112 | Perirhinal_area_35_L | 60 |
| 1113 | Perirhinal_area_36_L | 129 |
| 1114 | Medial_entorhinal_cortex_L | 272 |
| 1115 | Lateral_entorhinal_cortex_L | 51 |
| 1125 | 4th_ventricle_L | 94 |
| 1141 | lateral_lemniscus_unspecified_L | 47 |
| 1142 | Inferior_colliculus_dorsal_cortex_L | 44 |
| 1143 | Inferior_colliculus_central_nucleus_L | 105 |
| 1145 | Inferior_colliculus_external_cortex_L | 221 |
| 1146 | inferior_colliculus_brachium_L | 23 |
| 1150 | Medial_geniculate_body_marginal_zone_L | 12 |
| 1151 | Primary_auditory_area_L | 165 |
| 1152 | Secondary_auditory_area_dorsal_part_L | 148 |
| 1153 | Secondary_auditory_area_ventral_part_L | 127 |
| 1157 | external_medullary_lamina_auditory_radiation_L | 10 |
| 1180 | lateral_olfactory_tract_L | 43 |
| 1181 | Piriform_cortex_layer_1_L | 177 |
| 1182 | Piriform_cortex_layer_2_L | 55 |
| 1183 | Piriform_cortex_layer_3_L | 231 |
| 1184 | Nucleus_accumbens_core_L | 64 |
| 1187 | Substantia_nigra_reticular_part_L | 95 |
| 1188 | Substantia_nigra_compact_part_L | 19 |
| 1192 | Nucleus_accumbens_shell_L | 107 |
| 1193 | Ventral_pallidum_L | 84 |
| 1195 | Globus_pallidus_external_medial_part_L | 25 |
| 1196 | Ventral_tegmental_area_L | 35 |
| 1197 | Caudate_putamen_L | 1038 |
| 1198 | Globus_pallidus_external_lateral_part_L | 128 |
| 1199 | Ventral_striatal_region_unspecified_L | 37 |
| 1200 | Reticular_prethalamic_nucleus_unspecified_L | 106 |
| 1204 | Pregeniculate_nucleus_L | 21 |
| 1205 | Dorsal_lateral_geniculate_nucleus_L | 39 |
| 1208 | Posterior_intralaminar_nucleus_L | 11 |
| 1210 | Posterior_thalamic_nuclear_group_triangular_part_L | 11 |
| 1214 | Anteroventral_thalamic_nucleus_dorsomedial_part_L | 12 |
| 1215 | Anteroventral_thalamic_nucleus_ventrolateral_part_L | 23 |
| 1219 | Reuniens_thalamic_nucleus_L | 45 |
| 1221 | Ventromedial_thalamic_nucleus_L | 51 |
| 1227 | Ventral_posteromedial_thalamic_nucleus_L | 78 |
| 1228 | Laterodorsal_thalamic_nucleus_dorsomedial_part_L | 10 |
| 1229 | Laterodorsal_thalamic_nucleus_ventrolateral_part_L | 51 |
| 1230 | Posterior_thalamic_nucleus_L | 59 |
| 1231 | Ventrolateral_thalamic_nucleus_L | 60 |
| 1232 | Mediodorsal_thalamic_nucleus_lateral_part_L | 11 |
| 1233 | Mediodorsal_thalamic_nucleus_central_part_L | 13 |
| 1235 | Zona_incerta_dorsal_part_L | 36 |
| 1236 | Zona_incerta_ventral_part_L | 30 |
| 1240 | Mediodorsal_thalamic_nucleus_medial_part_L | 19 |
| 1242 | Paraventricular_thalamic_nuclei_anterior_and_posterior_L | 22 |
| 1246 | Paracentral_thalamic_nucleus_L | 22 |
| 1247 | Central_medial_thalamic_nucleus_L | 14 |
| 1249 | external_medullary_lamina_unspecified_L | 57 |
| 1254 | Anteromedial_thalamic_nucleus_L | 31 |
| 1267 | Parafascicular_thalamic_nucleus_L | 10 |
| 1270 | superior_cerebellar_peduncle_and_prerubral_field_L | 15 |
| 1281 | Subgeniculate_nucleus_L | 14 |
| 1283 | Lateral_posterior_thalamic_nucleus_lateral_part_L | 12 |
| 1285 | Lateral_posterior_thalamic_nucleus_mediorostral_part_L | 22 |
| 1287 | Zona_incerta_caudal_part_L | 23 |
| 1293 | Ventral_anterior_thalamic_nucleus_L | 10 |
| 1294 | Ventral_posterolateral_thalamic_nucleus_L | 75 |
| 1295 | Medial_geniculate_body_dorsal_division_L | 15 |
| 1298 | Medial_geniculate_body_ventral_division_L | 30 |
| 1400 | Ventrolateral_orbital_area_L | 64 |
| 1401 | Lateral_orbital_area_L | 117 |
| 1402 | Ventral_orbital_area_L | 65 |
| 1403 | Medial_orbital_area_L | 39 |
| 1404 | Dorsolateral_orbital_area_L | 92 |
| 1405 | Prelimbic_area_L | 176 |
| 1406 | Secondary_motor_area_L | 658 |
| 1407 | Frontal_association_area_3_L | 117 |
| 1408 | Primary_motor_area_L | 574 |
| 1409 | Agranular_insular_cortex_ventral_area_L | 51 |
| 1410 | Agranular_insular_cortex_dorsal_area_L | 72 |
| 1411 | Cingulate_area_1_L | 138 |
| 1412 | Claustrum_L | 63 |
| 1413 | Infralimbic_area_L | 34 |
| 1414 | Dysgranular_insular_cortex_L | 129 |
| 1416 | Granular_insular_cortex_L | 127 |
| 1417 | Primary_somatosensory_area_forelimb_representation_L | 366 |
| 1418 | Primary_somatosensory_area_dysgranular_zone_L | 99 |
| 1420 | Primary_somatosensory_area_face_representation_L | 520 |
| 1422 | Secondary_somatosensory_area_L | 416 |
| 1423 | Primary_somatosensory_area_hindlimb_representation_L | 173 |
| 1424 | Agranular_insular_cortex_posterior_area_L | 56 |
| 1425 | Primary_somatosensory_area_barrel_field_L | 486 |
| 1427 | Retrosplenial_dysgranular_area_L | 264 |
| 1429 | Primary_somatosensory_area_trunk_representation_L | 198 |
| 1430 | Retrosplenial_granular_area_L | 250 |
| 1432 | Parietal_association_cortex_lateral_area_L | 46 |
| 1433 | Parietal_association_cortex_medial_area_L | 57 |
| 1436 | Parietal_association_cortex_posterior_area_L | 53 |
| 1442 | Primary_visual_area_L | 544 |
| 1443 | Secondary_visual_area_lateral_part_L | 188 |
| 1444 | Temporal_association_cortex_L | 152 |
| 1448 | Secondary_visual_area_medial_part_L | 306 |
| 1500 | Endopiriform_nucleus_L | 102 |
| 1501 | Amygdaloid_area_unspecified_L | 470 |
| 1502 | Nucleus_of_the_lateral_olfactory_tract_L | 30 |

| **ROIs excluded** | | | |
| --- | --- | --- | --- |
| **ROI** | **NAME** | **VOXELS** | **RESOLUTION** |
|  | 1. **<10 voxel in native lateralised atlas** |  |  |
| 1054 | commissural_stria_terminalis_L | **8** | 0.15 |
| 70 | Central_canal_R | **5** | 0.15 |
| 1070 | Central_canal_L | **0** | 0.15 |
| 1080 | habenular_commissure_L | **5** | 0.15 |
| 1084 | medial_lemniscus_decussation_L | **0** | 0.15 |
| 1085 | pyramidal_decussation_L | **4** | 0.15 |
|  | 2. **<10 voxel after moving to pop avg space** |  |  |
| 46 | commissure_of_the_superior_colliculus_R | **5** | 0.2 |
| 1046 | commissure_of_the_superior_colliculus_L | **2** | 0.2 |
| 54 | commissural_stria_terminalis_R | **1** | 0.2 |
| 1063 | posterior_commissure_L | **8** | 0.2 |
| 72 | ascending_fibers_of_the_facial_nerve_R | **6** | 0.2 |
| 1072 | ascending_fibers_of_the_facial_nerve_L | **6** | 0.2 |
| 80 | habenular_commissure_R | **4** | 0.2 |
| 81 | Nucleus_of_the_stria_medullaris_R | **5** | 0.2 |
| 1081 | Nucleus_of_the_stria_medullaris_L | **1** | 0.2 |
| 132 | Superior_paraolivary_nucleus_R | **7** | 0.2 |
| 1132 | Superior_paraolivary_nucleus_L | **8** | 0.2 |
| 133 | Medial_superior_olive_R | **3** | 0.2 |
| 1133 | Medial_superior_olive_L | **2** | 0.2 |
| 140 | lateral_lemniscus_commissure_R | **5** | 0.2 |
| 1140 | lateral_lemniscus_commissure_L | **7** | 0.2 |
| 201 | Peripeduncular_nucleus_R | **4** | 0.2 |
| 1201 | Peripeduncular_nucleus_L | **4** | 0.2 |
| 223 | Angular_thalamic_nucleus_R | **7** | 0.2 |
| 1223 | Angular_thalamic_nucleus_L | **8** | 0.2 |
| 238 | Zona_incerta_A13_dopamine_cells_R | **5** | 0.2 |
| 1238 | Zona_incerta_A13_dopamine_cells_L | **6** | 0.2 |
| 260 | Intermediodorsal_thalamic_nucleus_R | **9** | 0.2 |
| 268 | Retroreuniens_thalamic_nucleus_R | **3** | 0.2 |
| 1268 | Retroreuniens_thalamic_nucleus_L | **2** | 0.2 |
| 272 | Intergeniculate_leaflet_R | **8** | 0.2 |
| 282 | Ethmoid-Limitans_nucleus_R | **9** | 0.2 |
| 1282 | Ethmoid-Limitans_nucleus_L | **8** | 0.2 |
| 290 | intramedullary_thalamic_area_R | **2** | 0.2 |
| 1290 | intramedullary_thalamic_area_L | **3** | 0.2 |
| 299 | Medial_geniculate_body_suprageniculate_nucleus_R | **9** | 0.2 |
|  | 3. **<10 voxel after exlcuding area outside FOV** |  |  |
| 45 | Spinal_cord_R | **0** | 0.2 |
| 1045 | Spinal_cord_L | **8** | 0.2 |
|  | 4. **<10 voxel after resampling to func res** |  |  |
| 3 | Subthalamic_nucleus_R | **4** | 0.3 |
| 1003 | Subthalamic_nucleus_L | **3** | 0.3 |
| 6 | alveus_of_the_hippocampus_R | **4** | 0.3 |
| 1006 | alveus_of_the_hippocampus_L | **3** | 0.3 |
| 1035 | facial_nerve_unspecified_L | **8** | 0.3 |
| 37 | anterior_commissure_posterior_limb_R | **8** | 0.3 |
| 1037 | anterior_commissure_posterior_limb_L | **5** | 0.3 |
| 1038 | ventral_hippocampal_commissure_L | **6** | 0.3 |
| 53 | mammillotegmental_tract_R | **1** | 0.3 |
| 1053 | mammillotegmental_tract_L | **6** | 0.3 |
| 57 | genu_of_the_facial_nerve_R | **5** | 0.3 |
| 1057 | genu_of_the_facial_nerve_L | **4** | 0.3 |
| 60 | fasciculus_retroflexus_R | **6** | 0.3 |
| 1060 | fasciculus_retroflexus_L | **7** | 0.3 |
| 63 | posterior_commissure_R | **3** | 0.3 |
| 68 | brachium_of_the_superior_colliculus_R | **3** | 0.3 |
| 1068 | brachium_of_the_superior_colliculus_L | **4** | 0.3 |
| 73 | anterior_commissure_intrabulbar_part_R | **8** | 0.3 |
| 1073 | anterior_commissure_intrabulbar_part_L | **8** | 0.3 |
| 83 | supraoptic_decussation_R | **6** | 0.3 |
| 1083 | supraoptic_decussation_L | **4** | 0.3 |
| 84 | medial_lemniscus_decussation_R | **1** | 0.3 |
| 85 | pyramidal_decussation_R | **4** | 0.3 |
| 123 | Ventral_cochlear_nucleus_granule_cell_layer_R | **9** | 0.3 |
| 1123 | Ventral_cochlear_nucleus_granule_cell_layer_L | **9** | 0.3 |
| 126 | Dorsal_cochlear_nucleus_molecular_layer_R | **9** | 0.3 |
| 1126 | Dorsal_cochlear_nucleus_molecular_layer_L | **6** | 0.3 |
| 127 | Dorsal_cochlear_nucleus_fusiform_and_granule_layer_R | **6** | 0.3 |
| 1127 | Dorsal_cochlear_nucleus_fusiform_and_granule_layer_L | **2** | 0.3 |
| 129 | acoustic_striae_R | **3** | 0.3 |
| 1129 | acoustic_striae_L | **3** | 0.3 |
| 1131 | Nucleus_of_the_trapezoid_body_L | **8** | 0.3 |
| 134 | Lateral_superior_olive_R | **7** | 0.3 |
| 1134 | Lateral_superior_olive_L | **5** | 0.3 |
| 136 | Ventral_periolivary_nuclei_R | **3** | 0.3 |
| 1136 | Ventral_periolivary_nuclei_L | **4** | 0.3 |
| 1137 | Lateral_lemniscus_ventral_nucleus_L | **6** | 0.3 |
| 138 | Lateral_lemniscus_intermediate_nucleus_R | **6** | 0.3 |
| 1138 | Lateral_lemniscus_intermediate_nucleus_L | **9** | 0.3 |
| 139 | Lateral_lemniscus_dorsal_nucleus_R | **5** | 0.3 |
| 1139 | Lateral_lemniscus_dorsal_nucleus_L | **6** | 0.3 |
| 150 | Medial_geniculate_body_marginal_zone_R | **6** | 0.3 |
| 159 | Ventral_cochlear_nucleus_posterior_part_R | **8** | 0.3 |
| 1159 | Ventral_cochlear_nucleus_posterior_part_L | **8** | 0.3 |
| 160 | Ventral_cochlear_nucleus_cap_area_R | **7** | 0.3 |
| 1160 | Ventral_cochlear_nucleus_cap_area_L | **7** | 0.3 |
| 162 | Spiral_ganglion_R | **1** | 0.3 |
| 1162 | Spiral_ganglion_L | **4** | 0.3 |
| 163 | Nucleus_sagulum_R | **9** | 0.3 |
| 1163 | Nucleus_sagulum_L | **7** | 0.3 |
| 164 | Reticular_prethalamic_nucleus_auditory_segment_R | **5** | 0.3 |
| 1164 | Reticular_prethalamic_nucleus_auditory_segment_L | **9** | 0.3 |
| 189 | Substantia_nigra_lateral_part_R | **6** | 0.3 |
| 1189 | Substantia_nigra_lateral_part_L | **5** | 0.3 |
| 1206 | Lateral_habenular_nucleus_L | **8** | 0.3 |
| 207 | Medial_habenular_nucleus_R | **3** | 0.3 |
| 1207 | Medial_habenular_nucleus_L | **8** | 0.3 |
| 208 | Posterior_intralaminar_nucleus_R | **8** | 0.3 |
| 1211 | Parataenial_thalamic_nucleus_L | **5** | 0.3 |
| 213 | Anterodorsal_thalamic_nucleus_R | **7** | 0.3 |
| 1213 | Anterodorsal_thalamic_nucleus_L | **8** | 0.3 |
| 216 | Rhomboid_thalamic_nucleus_R | **4** | 0.3 |
| 1216 | Rhomboid_thalamic_nucleus_L | **3** | 0.3 |
| 218 | Xiphoid_thalamic_nucleus_R | **4** | 0.3 |
| 1218 | Xiphoid_thalamic_nucleus_L | **7** | 0.3 |
| 1222 | Submedius_thalamic_nucleus_L | **8** | 0.3 |
| 239 | pretectothalamic_lamina_R | **5** | 0.3 |
| 1239 | pretectothalamic_lamina_L | **3** | 0.3 |
| 1248 | Central_lateral_thalamic_nucleus_L | **7** | 0.3 |
| 255 | Interanteromedial_thalamic_nucleus_R | **6** | 0.3 |
| 1255 | Interanteromedial_thalamic_nucleus_L | **3** | 0.3 |
| 257 | Zona_incerta_rostral_part_R | **6** | 0.3 |
| 1257 | Zona_incerta_rostral_part_L | **7** | 0.3 |
| 1260 | Intermediodorsal_thalamic_nucleus_L | **2** | 0.3 |
| 266 | Ventral_posterior_nucleus_of_the_thalamus_parvicellular_part_R | **5** | 0.3 |
| 1266 | Ventral_posterior_nucleus_of_the_thalamus_parvicellular_part_L | **4** | 0.3 |
| 1272 | Intergeniculate_leaflet_L | **2** | 0.3 |
| 278 | Subparafascicular_nucleus_R | **4** | 0.3 |
| 1278 | Subparafascicular_nucleus_L | **7** | 0.3 |
| 280 | Fields_of_Forel_R | **3** | 0.3 |
| 1280 | Fields_of_Forel_L | **5** | 0.3 |
| 284 | Zona_incerta_A11_dopamine_cells_R | **4** | 0.3 |
| 1284 | Zona_incerta_A11_dopamine_cells_L | **2** | 0.3 |
| 286 | Lateral_posterior_thalamic_nucleus_mediocaudal_part_R | **9** | 0.3 |
| 1286 | Lateral_posterior_thalamic_nucleus_mediocaudal_part_L | **9** | 0.3 |
| 291 | internal_medullary_lamina_R | **4** | 0.3 |
| 1291 | internal_medullary_lamina_L | **6** | 0.3 |
| 297 | Medial_geniculate_body_medial_division_R | **5** | 0.3 |
| 1297 | Medial_geniculate_body_medial_division_L | **9** | 0.3 |
| 1299 | Medial_geniculate_body_suprageniculate_nucleus_L | **6** | 0.3 |
|  | 5. **<10 voxel after multipling by inclusive mask** |  |  |
| 35 | facial_nerve_unspecified_R | **0** | 0.3 |
| 41 | optic_nerve_R | **3** | 0.3 |
| 1041 | optic_nerve_L | **5** | 0.3 |
| 74 | Inferior_olive_R | **0** | 0.3 |
| 1074 | Inferior_olive_L | **0** | 0.3 |
| 75 | Spinal_trigeminal_nucleus_R | **2** | 0.3 |
| 1075 | Spinal_trigeminal_nucleus_L | **6** | 0.3 |
| 119 | Vestibular_apparatus_R | **0** | 0.3 |
| 1119 | Vestibular_apparatus_L | **0** | 0.3 |
| 120 | Cochlea_R | **0** | 0.3 |
| 1120 | Cochlea_L | **0** | 0.3 |
| 121 | Cochlear_nerve_R | **0** | 0.3 |
| 1121 | Cochlear_nerve_L | **0** | 0.3 |
| 122 | Vestibular_nerve_R | **4** | 0.3 |
| 1122 | Vestibular_nerve_L | **2** | 0.3 |
| 1128 | Dorsal_cochlear_nucleus_deep_core_L | **9** | 0.3 |
| 130 | trapezoid_body_R | **0** | 0.3 |
| 1130 | trapezoid_body_L | **0** | 0.3 |
| 131 | Nucleus_of_the_trapezoid_body_R | **0** | 0.3 |
| 135 | Superior_periolivary_region_R | **0** | 0.3 |
| 1135 | Superior_periolivary_region_L | **0** | 0.3 |
| 137 | Lateral_lemniscus_ventral_nucleus_R | **9** | 0.3 |
| 158 | Ventral_cochlear_nucleus_anterior_part_R | **2** | 0.3 |
| 1158 | Ventral_cochlear_nucleus_anterior_part_L | **1** | 0.3 |
